# Longitudinal analysis of a dietary mouse model of non-alcoholic fatty liver disease (NAFLD) and non-alcoholic steatohepatitis (NASH)

**DOI:** 10.1101/2023.05.19.540989

**Authors:** Marissa Saenz, Jillian C. McDonough, Elizabeth Bloom-Saldana, Jose M. Irimia, Emily L. Cauble, Ashly Castillo, Patrick T. Fueger, Lindsey S. Treviño

**Affiliations:** Center for Comparative Medicine; Department of Molecular and Cellular Endocrinology; Division of Health Inequities, Department of Population Sciences; Comprehensive Metabolic Phenotyping Core, Beckman Research Institute, City of Hope, Duarte, CA; Eugene and Ruth Roberts Summer Student Academy, Irell & Manella Graduate School of Biological Sciences, City of Hope, Duarte, CA; Cancer Control and Population Sciences, Comprehensive Cancer Center, City of Hope, Duarte, CA

**Author notes:** **To whom correspondence should be addressed:** Lindsey S. Treviño Patrick T. Fueger Beckman Research Institute of City of Hope 1500 E Duarte Rd Duarte, CA 91010.

**Keywords:** microbiome, liver, metabolism, transcriptome, steatosis

## Abstract

Non-alcoholic fatty liver disease (NAFLD), and resultant non-alcoholic steatohepatitis (NASH), incidence and prevalence are rising globally due to increasing rates of obesity and diabetes. Currently, there are no approved pharmacological treatments for NAFLD, highlighting a need for additional mechanistic studies to develop prevention and/or therapeutic strategies. Diet-induced preclinical models of NAFLD can be used to examine the dynamic changes that occur during NAFLD development and progression throughout the lifespan. To date, most studies utilizing such models have focused exclusively on terminal time points and have likely missed critical early and late changes that are important for NAFLD progression (i.e, worsening). We performed a longitudinal analysis of histopathological, biochemical, transcriptomic, and microbiome changes that occurred in adult male mice fed either a control diet or a NASH-promoting diet (high in fat, fructose, and cholesterol) for up to 30 weeks. We observed progressive development of NAFLD in mice fed the NASH diet compared to the control diet. Differential expression of immune-related genes was observed at an early stage of diet-induced NAFLD development (10 weeks) and persisted into the later stages of the disease (20 and 30 weeks). Differential expression of xenobiotic metabolism related genes was observed at the late stage of diet-induced NAFLD development (30 weeks). Microbiome analysis revealed an increased abundance of *Bacteroides* at an early stage (10 weeks) that persisted into the later stages of the disease (20 and 30 weeks). These data provide insight into the progressive changes that occur during NAFLD/NASH development and progression in the context of a typical Western diet. Furthermore, these data are consistent with what has been reported in patients with NAFLD/NASH, supporting the preclinical use of this diet-induced model for development of strategies to prevent or treat the disease.

---

Recent estimates suggest that one in four people have non-alcoholic fatty liver disease (NAFLD), making NAFLD the most common liver disease (1, 2). The global rise in obesity, which promotes metabolic dysfunction is thought to be fueling this rise in NAFLD. Thus, a recent consensus has suggested a switch to the term metabolic (dysfunction) associated fatty liver disease (MAFLD) (3). Whereas NALFD and MAFLD themselves could be relatively benign, they are precursors for the development of severe liver diseases, including non-alcoholic steatohepatitis (NASH), hepatic cirrhosis, and hepatocellular carcinoma (HCC). One report suggested that cases of NASH will increase 63% from 2015 to 2030, going from 16.5 to 27.0 million cases (4). Unfortunately, despite a quite dense therapeutic pipeline, there are currently no approved therapies for treating NAFLD and NASH other than liver transplantation.

Because of the explosion of cases of NAFLD and MAFLD worldwide and the potential for surging numbers of patients with more severe liver diseases, there is great urgency to develop new therapeutics to combat this spectrum of diseases. To this end, investigators commonly employ animal models for both discovery work into the basic biological mechanisms of disease pathogenesis as well as therapeutic development and validation. Historically, mouse models of NAFLD, NASH, and HCC have relied on genetic manipulation, such as those modulating feeding behavior (e.g., *ob/ob* mice, *db/db* mice, *Mcr4^-/-^* mice) or the use of carcinogens such as diethylnitrosamine (DEN), which is often coupled with methionine- and choline-deficient diets or those laden with lipids (reviewed in (5)). However, the use of exclusively nutritionally-based animal models has been less common. Further, most studies have focused solely on terminal time points and have not examined the dynamic changes throughout the lifespan, thereby likely missing critical changes occurring during the prolonged exposure to the diet and potential intervention points during the progression of the disease. Therefore, we aimed to longitudinally characterize an animal model of MALFD and NASH reliant solely on dietary manipulation with one consisting of elevated fat, cholesterol, and fructose. The emphasis on a longitudinal study was essential to highlight time-dependent changes in response to the diet. This approach offers key insights into predictive changes in the microbiome and hepatic transcriptome and pathology that can inform subsequent disease progression.

By comparing mice fed either a control or NASH-promoting diet over time, we examined a model that closely recapitulated several features of human NASH, including histological, transcriptional, and biochemical parameters in the liver as well as the microbiome. In addition, we were able to track changes in parameters over the course of the study to establish a sequence of events related to the pathogenesis of NASH. This model can be used for both mechanistic studies or therapeutic development for the prevention or treatment of NAFLD, MAFLD, and NASH.

## MATERIALS AND METHODS

### Animal studies

All animals were handled and cared for according to protocols approved by the City of Hope Institutional Animal Care and Use Committee. Mice were maintained on a standard 12-h light-dark cycle and had free access to diet and water. Six-week-old male C57Bl/6J mice were fed either a Western-style diet (NASH, 40% kcal from fat, Research Diets D09100310) or a control diet (CON, 10% kcal from fat, Research Diets D09100304) for up to 30 weeks (**Figure 1A**). Mice were weighed weekly and body composition was assessed at 10, 20, and 30 weeks by quantitative magnetic resonance technology (EchoMRI™ 3-in-1, Houston, TX). Glucose tolerance and insulin tolerance tests were conducted at approximately 8, 18, and 28 weeks on the diet and were separated by at least three days. For glucose tolerance testing (GTT), 1.5 g/kg body weight of D-glucose (Sigma-Aldrich, St. Louis, MO) was administered by oral gavage into 6-h fasted mice. For insulin tolerance testing (ITT), 0.75 U/kg body weight of Humulin (Eli Lilly & Co., Indianapolis, IN) was injected intraperitoneally into 6-h fasted mice. Blood was sampled from a tail vein at the indicated time points, and blood glucose was measured using an AlphaTrak glucometer (Abbott Laboratories, Abbott Park, IL). A week following tolerance testing, blood was withdrawn from a submandibular vein for measuring hormones and analytes in the plasma using commercial kits. After harvesting, blood was centrifuged at 12,500 x *g*, plasma was collected, and stored at -80 until analysis. At 9, 19, and 29 weeks on the diet, mice were placed individually into instrumented metabolic cages to measure gas exchange, energy expenditure, food/water consumption, and activity (Promethion, Sable Systems). Separate cohorts were terminated after 10, 20, or 30 weeks on the respective diets. Livers were immediately harvested, weighed, and flash-frozen in liquid nitrogen for follow-up studies.

**Figure 1.**
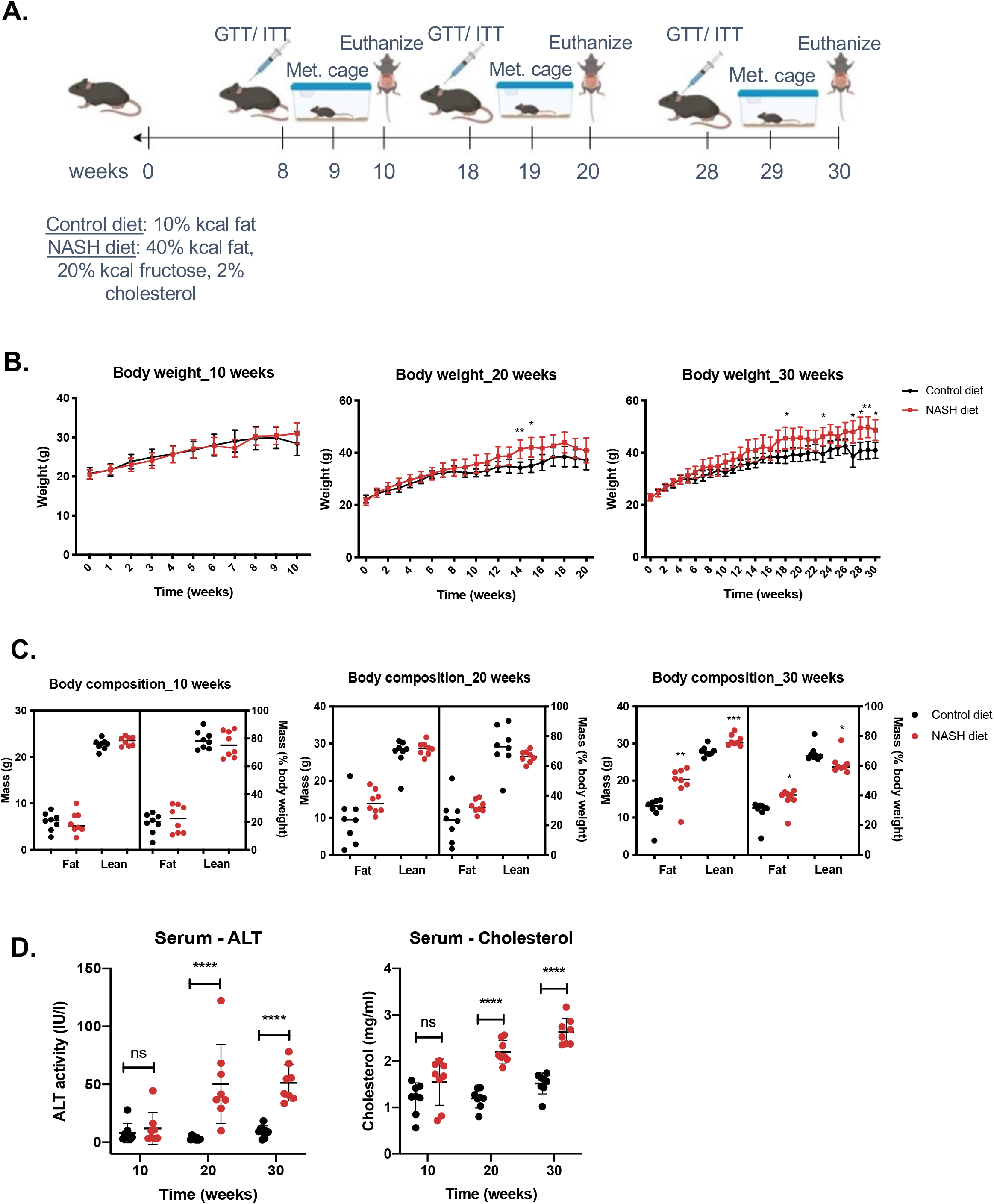
Characterization of mice fed a control or NASH diet for up to 30 weeks. Schematic of experimental design (A). Effect of feeding male mice with either a control diet (10% kcal from fat) or NASH diet (40% kcal from fat, 20% kcal fructose, and 2% cholesterol) on body weight (B) and body composition (C) for up to 30 weeks. Effect of the diets on serum levels of alanine aminotransferase (ALT) and cholesterol (D). Significant differences in body weight were determined by ANOVA. Significant differences in body composition and serum levels of ALT and cholesterol were determined using Student’s t-tests. Changes were considered significant at p < 0.05. * p < 0.05, ** p < 0.01, **** p<0.0001.

### Blood analysis

All blood analysis was conducted by the Comprehensive Metabolic Phenotyping Core at City of Hope. Commercial assays performed according to the manufacturers’ protocols and included: alanine transaminase (ALT; #A7526-01-1953, Pointe Scientific), total cholesterol (#MAK043, Sigma), albumin (#MAK125, Sigma), insulin (#SRI-13K, Millipore), free glycerol and triglycerides (#TR0100, Sigma), and non-esterified free fatty acids (#NEFA-HR, Wako).

### Liver pathology

At harvest, liver weights were recorded, and samples were either flash-frozen or formalin-fixed for downstream assays. Hepatic glycogen content was determined by measuring amyloglucosidase-released glucose from glycogen as previously described (6). For measuring hepatic triglyceride content, the lipid fraction of the liver was extracted as previously described (7), and total triglycerides were measured enzymatically using a triglyceride and free glycerol determination kit (#TR0100, Sigma) following the manufacturer’s instructions.

Formalin-fixed liver tissue was paraffin embedded at the Research Pathology Core at City of Hope. Blocks were submitted to HistoWiz for hematoxylin and eosin (H&E) and Masson’s Trichrome staining. Stained slides were reviewed by an independent, blinded pathologist contracted by HistoWiz and evaluated for degree of fatty change/fat accumulation (steatosis), lobular inflammation, and fibrosis. Steatosis scores were assigned as follows: <5% = 0, 33% = 1, 33-60% = 2, >60% = 3. Scores for lobular inflammation were assigned as follows: none = 0, <2 sites = 1, 2-4 sites = 3, >4 sites = 4. Fibrosis scores were assigned as follows: none = 0, minimal = 1, mild = 2, moderate = 3, severe = 4.

### RNA-sequencing analysis of whole liver

Frozen liver tissue (n = 3 per timepoint per diet group) was sent to Active Motif for RNA extraction, cDNA generation, library prep, and RNA-sequencing (RNA-seq) analysis (sequenced as 42-bp paired-end reads). Briefly, total RNA was isolated using the Qiagen RNeasy Mini Kit per the manufacturer’s instructions (#74104, Qiagen). Total RNA was then used in Illumina’s TruSeq Stranded mRNA Library kit (#20020594). Libraries were sequenced on Illumina NextSeq 550 as paired-end 42-nt reads. Sequence reads were analyzed with the STAR alignment – DESeq2 software pipeline by mapping to the mm10 assembly genome using the STAR 2.5.2b aligner software, followed by fragment assignment to annotated genes. Differential analysis was performed to identify differentially expressed genes (DEGs) using DESeq2 at 0.1 FDR. Using DESeq2 normalized gene counts, gene set enrichment analysis (GSEA) was performed to determine whether members of *a priori* defined gene sets (within MSigDBs C5 GO gene set) are enriched. Principle component analysis (PCA) was also performed to determine how the different diets can account for the variation across samples. These data analyses were performed at Active Motif. In addition, we performed Ingenuity Pathway Analysis (IPA) to identify signaling pathways enriched in the identified DEGs, as well as to identify potential upstream regulators of the DEGs.

### Real-time quantitative PCR

Frozen liver samples (n = 8 per timepoint per diet group) were crushed with a tissue pulverizer before total RNA was extracted from using the Qiagen RNeasy Mini Kit (Qiagen, Germantown, MD) per manufacturer’s instructions. RNA concentration was measured using a NanoDrop One (Thermo Fisher Scientific, Waltham, MA). A total of 500 ng of RNA was converted to cDNA using the High-Capacity RNA-to-cDNA Kit (Applied Biosystems, Thermo Fisher Scientific, Waltham, MA). The PowerUp SYBR Green Master Mix (Applied Biosystems, Thermo Fisher Scientific, Waltham, MA), plus primer pairs **(Table 1)** were used to validate DEGs in pathways of interest.

**Table 1:**
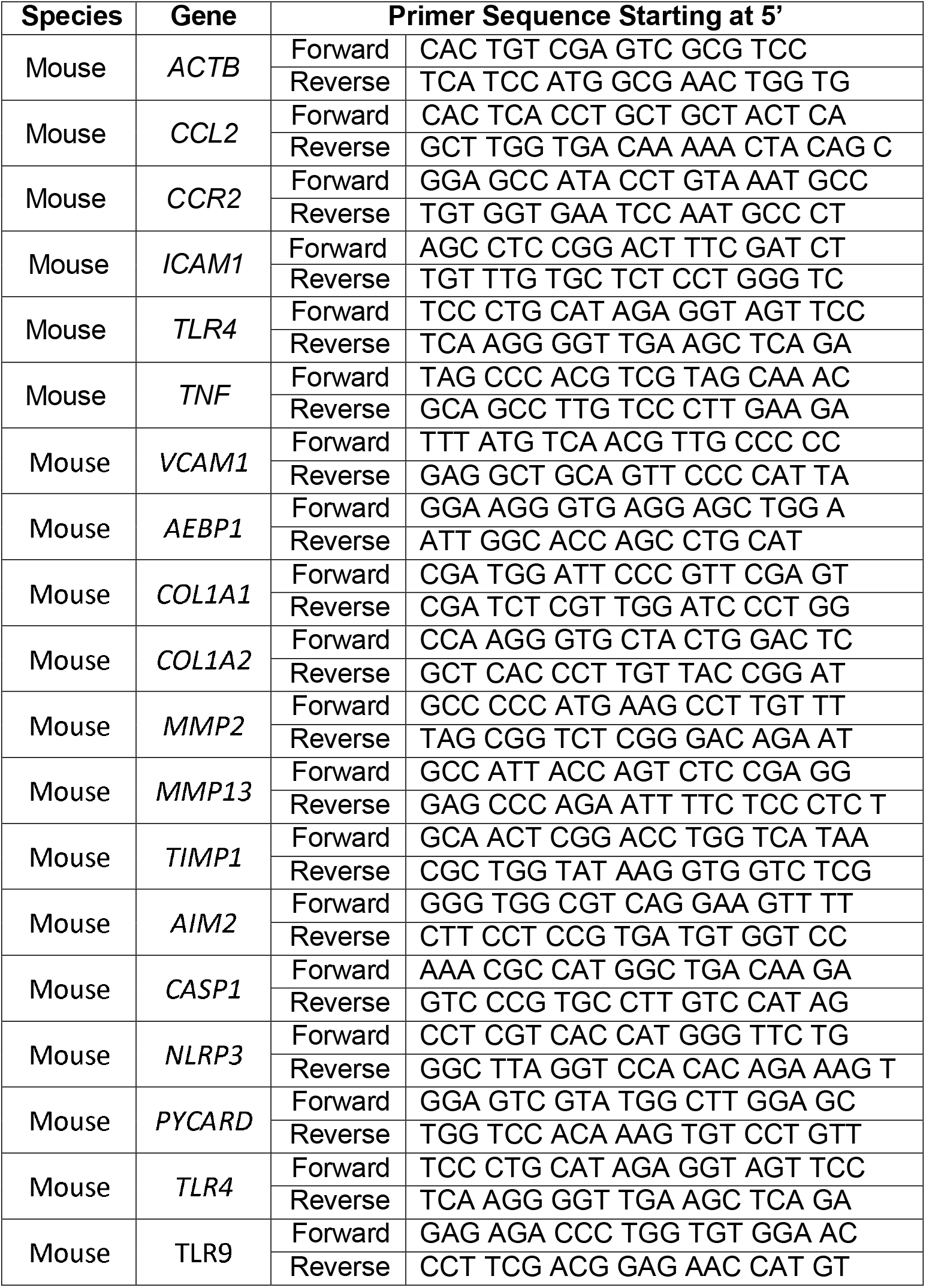
Oligonucleotide primer sequences for real-time quantitative PCR.

### Fecal microbiome analysis by whole-genome sequencing (WGS)

Fecal samples were collected from mice while they were individually housed in metabolic cages and then flash-frozen for downstream analysis. Frozen samples (2 per mouse at each time point) were immersed in the Transnetyx microbiome sample collection tube (1.5 mL, prefilled with DNA stabilization liquid and glass beads) and sent to Transnetyx (Cordova, TN) for automated DNA extraction (Qiagen DNeasy 96 PowerSoil Pro QIAcube HT extraction kit), KAPA HyperPlus library prep, and shallow shotgun whole genome sequencing (WGS; minimum read depth of 2 million paired-end reads) on an Illumina NovaSeq instrument. Shallow WGS provides species/sub-strain level taxonomic resolution, as well as identification of non-bacterial organisms that may be present in the sample. Data analysis and visualization of taxonomy, alpha-diversity, beta-diversity, and PCA clustering were performed using One Codex (http://www.onecodex.com). The One Codex Database contains >115,000 completed microbial reference genomes, including 62,000 distinct bacterial genomes; human and mouse genomes were included to exclude host reads. Individual sequences were compared by exact alignment using k-mers, where k=31, as previously described (8-10). Relative species abundance was estimated from the sequencing depth and coverage across the reference genomes.

### Statistical analysis

All experiments were performed at least three independent times. Where appropriate, significance was determined by one- or two-tailed Student’s *t*-test or one- or two-way ANOVA with Bonferroni multiple comparison tests. Metabolic cage data were analyzed by ANCOVA in CalR with weight as a covariate (11). Differences were considered significant when *P* < 0.05. Data are reported as means ± SEM.

## RESULTS

### Impact of NASH diet on metabolic parameters

Male mice fed the NASH diet exhibited increased body weight after 27 weeks compared to mice fed the control diet **(Figure 1B)**. Although there was a significant increase in body weight in the mice fed the NASH diet at weeks 14 and 15 for the 20-week cohort and at weeks 18 and 23 for the 30-week cohort, this increase was not sustained throughout the course of the study. To quantify body composition changes we determined both the absolute and relative fat and lean masses using EchoMRI. By 30 weeks on the NASH diet, both absolute and relative fat mass were increased, and whereas absolute lean mass was increased, relative lean mass was decreased due the dramatic increase in absolute fat mass **(Figure 1C)**. Serum alanine transaminase (ALT) levels were measured as a marker of liver damage. By 20 weeks on the NASH diet, serum ALT levels increased and remained elevated at 30 weeks on the diet **(Figure 1D)**. Similarly, serum cholesterol levels were also significantly elevated in the NASH diet-fed animals at 20 and 30 weeks **(Figure 1D)**. There were no significant differences between the two groups in serum ALT or cholesterol at the 10-week time point **(Figure 1D)** or serum albumin, insulin, free glycerol, triglycerides, or non-esterified fatty acids (NEFAs) throughout the study **(Supplemental Figure 1)**. We also performed oral glucose (GTT) and intraperitoneal insulin tolerance (ITT) tests at 10, 20, and 30 weeks on the respective diets. Surprisingly, we did not detect a significant difference in either GTT or ITT at any time point between the two diet groups, which could be due to the relatively mild changes in adiposity in the current dietary model **(Supplemental Table 1)**.

Metabolic caging was used to assess gas exchange, food/water consumption, activity, and energy expenditure **(Figure 2 and Supplemental Figures 2 and 3)**. Daily food consumption on a per kcal basis as well as food consumption during both dark and light phases was greater in the NASH diet-fed animals compared to control diet-fed animals after 10 weeks on the diet, yet body weight and composition were unaltered at this point in the study. There were no discernable differences in gas exchange, activity, energy expenditure, energy balance, or respiratory exchange ratio (RER) between groups at the 10- and 20-week time points. At 30 weeks, RER was decreased throughout the day as well as both the light and dark cycles in the NASH diet-fed group compared to the control diet-fed group, suggesting a preference for fat utilization over carbohydrate utilization. Other metabolic cage parameters were not different between groups at the 30-week time point.

**Figure 2.**
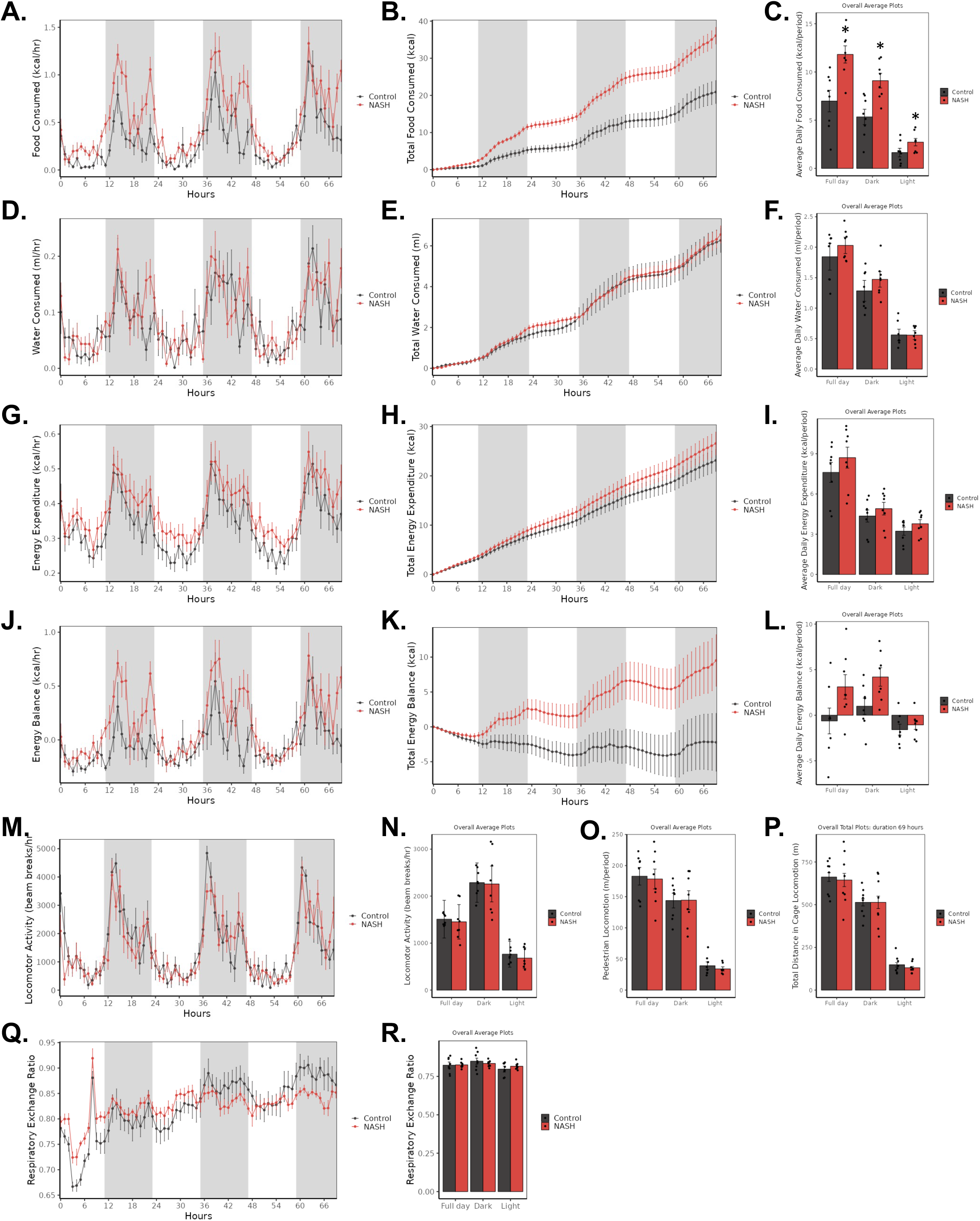
Increased food consumption in NASH diet-fed mice. Instrumented metabolic caging was used to quantify food consumption (A-C), water consumption (D-F), energy expenditure (G-I), net energy balance (J-L), locomotor activity (M-P), and respiratory exchange ratio (Q-R) over a three-day period in mice fed a control or NASH diet for 10 weeks. n = 8; * p < 0.05.

### Progressive development of NAFLD in mice fed the NASH diet

Male mice fed the NASH diet for 10, 20, and 30 weeks exhibited significantly increased liver weight (measured as a percentage of body weight) **(Figure 3A)**. Whereas hepatic glycogen was significantly decreased at 10, 20, and 30 weeks on the diet, hepatic triglycerides were significantly increased at all three time points **(Figure 3B)**, suggesting a preferential shift in fuel source utilization versus storage. Corroborating the hepatic triglyceride data, mice fed the NASH diet exhibited histological accumulation of lipids in the liver (steatosis), as well as liver scarring (fibrosis) as early as 10 weeks on the diet **(Figure 3C)**. It should be noted that age-dependent liver steatosis was also observed in the mice fed the control diet, but not to the same extent of severity observed in the mice fed the NASH diet. For example, 75% (6/8) of the mice fed the NASH diet exhibited fat accumulation in >60% of the liver at 30 weeks (as determined by pathology scoring), whereas none (0/8) of the mice fed the control diet were assigned this score **(Figure 3D)**. Similar results were observed for liver fibrosis, with 63% (5/8) of the NASH diet fed mice and none (0/8) of the control fed diet mice scoring either mild, moderate, or severe for fibrosis at 30 weeks **(Figure 3D)**. As expected, the severity of steatosis and fibrosis progressively worsened over time in the mice fed the NASH diet **(Figure 3D)**. There was no significant difference in lobular inflammation between the two diet groups (data not shown).

**Figure 3.**
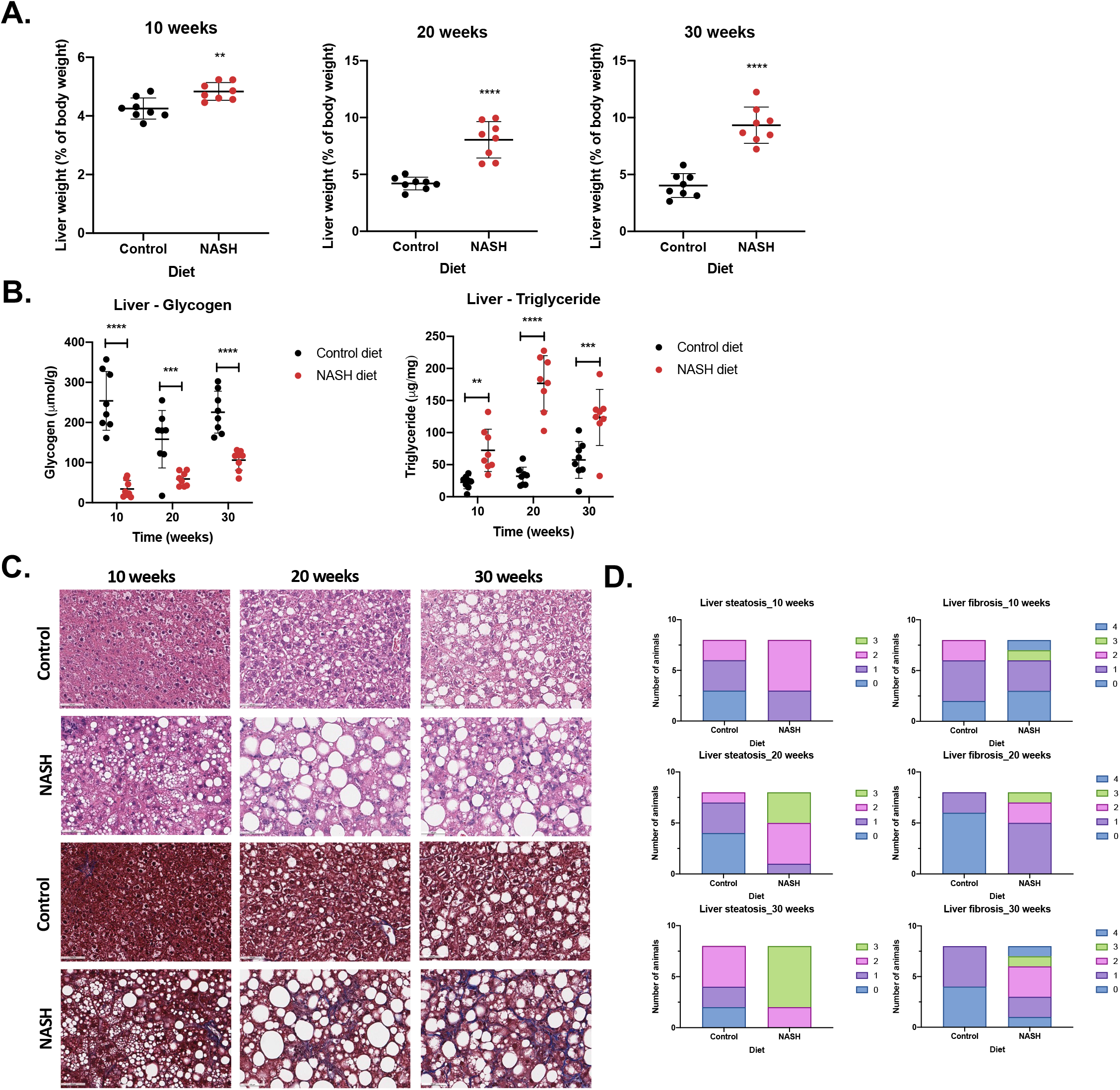
Biochemical and histological analysis of livers from mice fed a NASH diet for up to 30 weeks. Effect of feeding male mice with either a control diet (10% kcal from fat) or NASH diet (40% kcal from fat, 20% kcal fructose, and 2% cholesterol) on liver weight (as a percent of body weight) (A), as well as on levels of glycogen and triglycerides in the liver (B) at 10, 20, and 30 weeks on the respective diets. Representative hematoxylin and eosin (H&E) and Masson’s Trichrome stained sections of liver harvested after 10, 20, or 30 weeks on a control or NASH diet (C). Scale bar = 100 μM. Bar graphs summarizing steatosis and fibrosis scores after 10, 20, or 30 weeks on a control or NASH diet. Scores were determined by an independent, blinded pathologist contracted by HistoWiz. Significant differences in liver weight, as well as liver glycogen and triglycerides were determined by Student’s t-tests at individual time points. Changes were considered significant at p < 0.05. ** p < 0.01, *** p < 0.001, **** p < 0.0001.

### Activated immune response in the liver of mice fed the NASH diet for 10 weeks

To determine changes in gene expression that accompany the changes in pathology observed in the mice fed the NASH diet, we performed RNA-sequencing with liver samples from mice fed the control or NASH diet for 10, 20, or 30 weeks. Principal component analysis (PCA) of the sequencing data revealed that 75% (61.86% PC1 and 13.1% PC2) of the variance between samples can be explained by diet **(Supplemental Figure 4)**. Approximately 71% (52.01% PC1 and 19.11% PC2) of the variance can be explained by both diet and time point **(Supplemental Figure 4)**. At 10 weeks, there were a total of 3,455 differentially expressed genes (DEGs; 2,155 upregulated and 1,300 downregulated) in the liver of mice fed the NASH diet compared to mice fed a control diet. The top 40 DEGs at 10 weeks included genes related to lipid metabolism and cholesterol biosynthesis **(Figure 4A)**. Ingenuity Pathway Analysis (IPA) revealed that the DEGs at this timepoint were enriched for pathways related to immune signaling, including, but not limited to, activation of TREM1 signaling, the Th1 pathway, and the Neuroinflammation Signaling Pathway **(Figure 4B)**. Using IPA, we also identified potential upstream regulators of the DEGs. These upstream regulators included mediators of immune responses such as interferon gamma, interleukin-2, interleukin-4, and tumor necrosis factor **(Figure 4C)**. Using Gene Set Enrichment Analysis (GSEA), we determined that the gene signature identified in liver from mice fed the NASH diet for 10 weeks is enriched for genes related to the adaptive immune response **(Figure 4D)**. We validated the expression of a subset of adaptive immune response-related genes in the larger cohort of animals, and observed increased expression of *Ccl2*, *Ccr2*, *Icam1*, *Tlr4*, *Tnf*, and *Vcam1* in the liver of mice fed the NASH diet for 10 weeks compared to the liver of mice fed the control diet **(Figure 4E)**.

**Figure 4.**
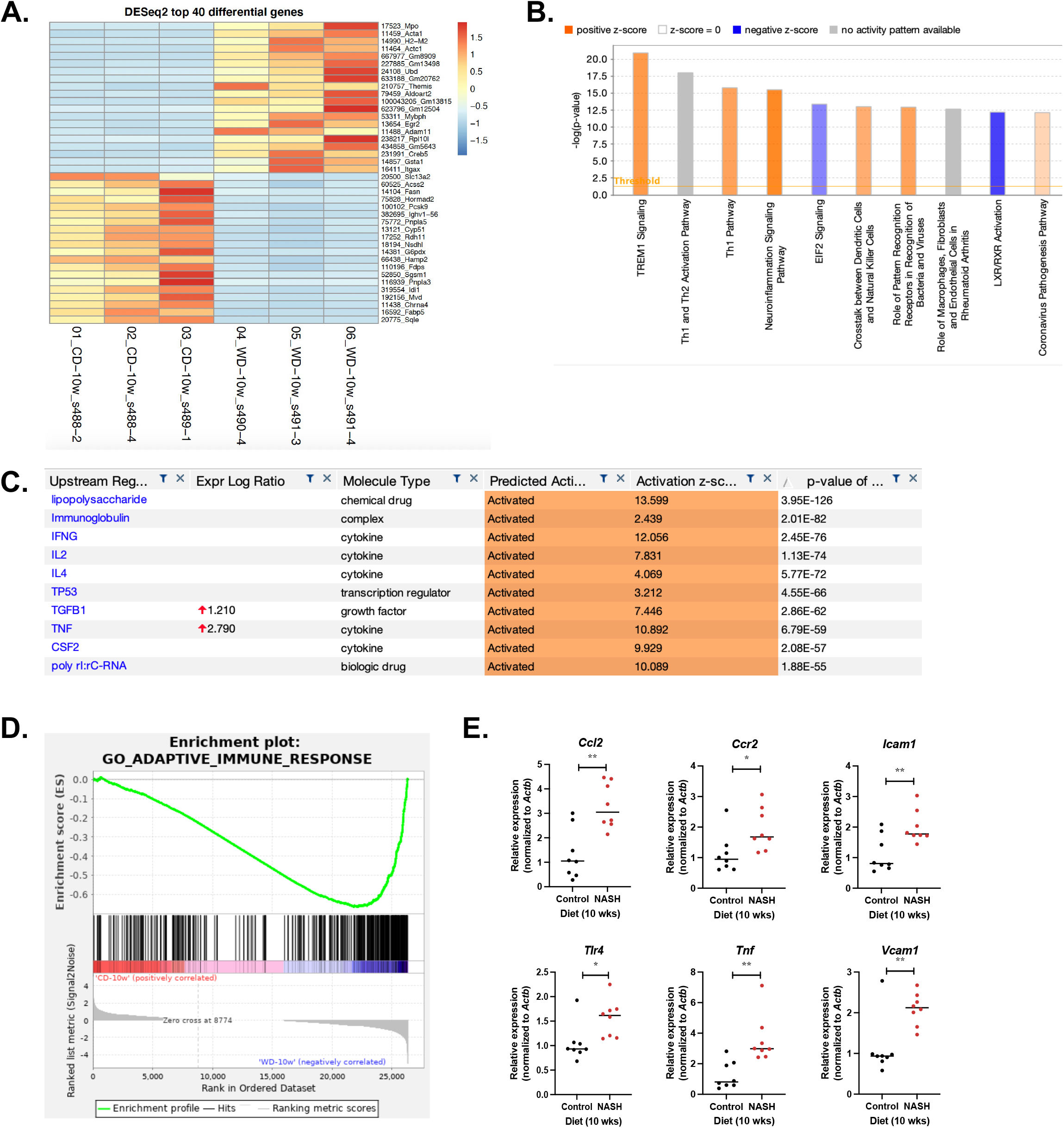
Hepatic transcriptome from mice fed a NASH diet for 10 weeks. Effect of feeding male mice either a control diet (10% kcal from fat) or NASH diet (40% kcal from fat, 20% kcal fructose, and 2% cholesterol) for 10 weeks on hepatic gene expression. Heat map of the top 40 differentially expressed genes (DEGs; A). Pathways related to immune response, including TREM1 Signaling, Th1 Pathway, and Neuroinflammation Signaling Pathway, are activated in the liver of animals fed a Western-style diet for 10 weeks compared to a control diet (B). Orange bars denote activated pathways (positive z-score > 2), whereas blue bars denote inhibited pathways (negative z-score <- 2.0). Putative upstream regulators of the DEGs at this time point include immune-related molecules such as interferon gamma, interleukin 2, interleukin 4, and tumor necrosis factor (C). In the liver of mice fed a NASH diet, the gene set for adaptive immune response is significantly enriched (D). Genes related to the adaptive immune response, including *Ccl2*, *Ccr2*, *Icam1*, *Tlr4*, *Tnf*, and *Vcam1* are significantly upregulated in the liver of mice fed the NASH diet for 10 weeks, compared to the control diet (E). Statistical significance was determined by t-test, with p < 0.05 considered significant. * p < 0.05, ** p <0.01.

### Activated fibrotic response in the liver of mice fed the NASH diet for 20 weeks

At 20 weeks, there were a total of 4,070 DEGs (2,286 upregulated and 1,784 downregulated) in the liver of mice fed the NASH diet compared to mice fed a control diet. The top 40 DEGs at 20 weeks included genes related to cytochrome P450 activity, secondary metabolite biosynthesis, transport, and catabolism, as well as genes that encode glycoproteins **(Figure 5A)**. Using IPA, we determined that DEGs at this time point were enriched for pathways related to fibrosis, including activation of the Hepatic Fibrosis Signaling Pathway and Pulmonary Fibrosis Idiopathic Signaling Pathway **(Figure 5B)**. Putative upstream regulators of the 20-week DEGs included fibrosis-related genes, such as *TGFB-1* (the master regulator of fibrosis) and *AGT* **(Figure 5C)**. Using GSEA, we determined that the gene signature identified in the liver from mice fed the NASH diet for 20 weeks was enriched for genes related to the GO term “Collagen Fibril Organization” **(Figure 5D)**. We validated a subset of fibrosis-related genes in the larger cohort of animals, and observed increased expression of *Aebp1*, *Col1a1*, *Col1a2*, *Mmp2*, *Mmp13*, and *Timp1* in the liver of mice fed the NASH diet for 20 weeks compared to the liver of mice fed the control diet **(Figure 5E)**.

**Figure 5.**
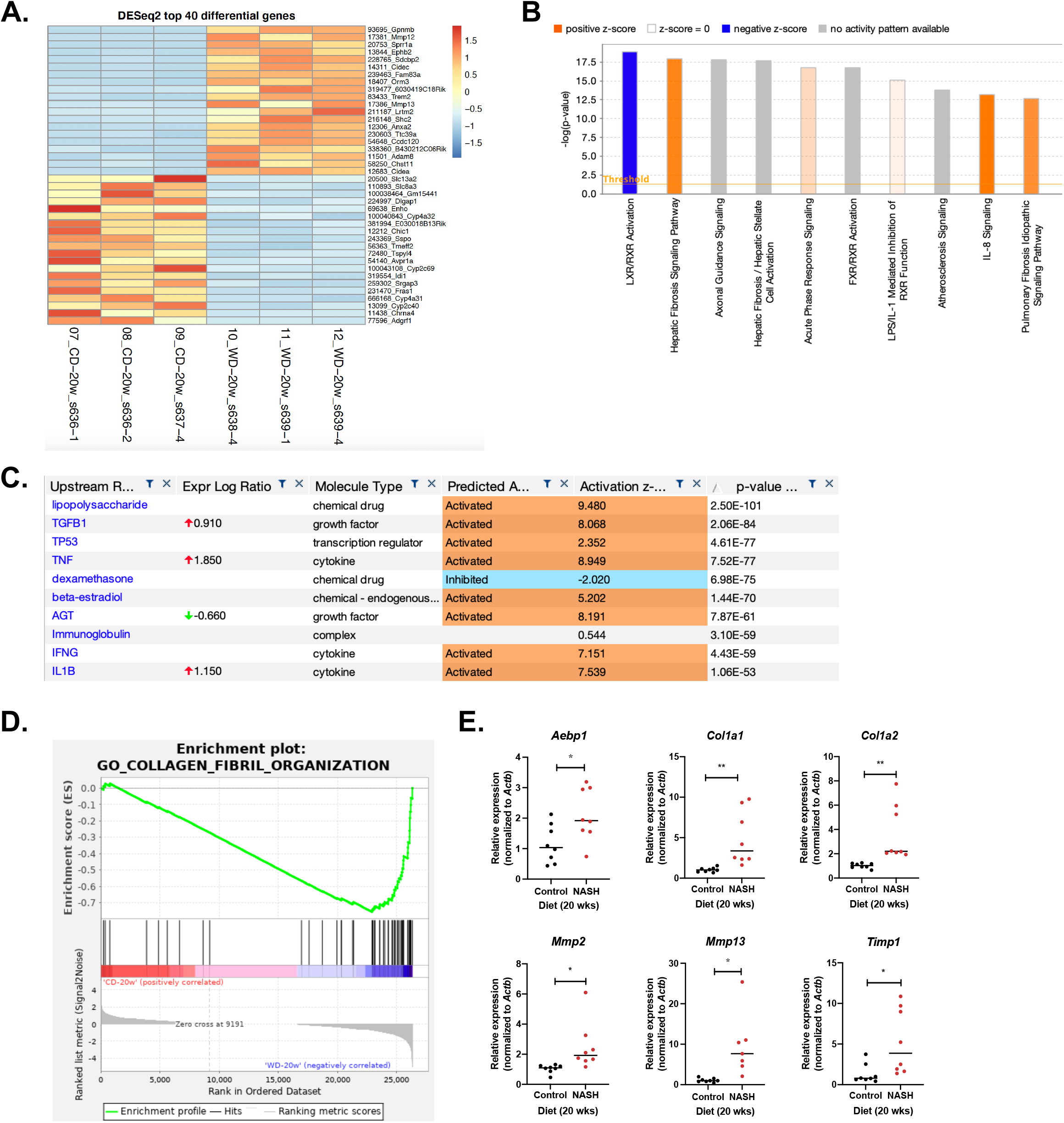
Hepatic transcriptome from mice fed a NASH diet for 20 weeks. Effect of feeding male mice either a control diet (10% kcal from fat) or NASH diet (40% kcal from fat, 20% kcal fructose, and 2% cholesterol) for 20 weeks on hepatic gene expression. Heat map of the top 40 differentially expressed genes (DEGs; A). Pathways related to fibrosis, including Hepatic Fibrosis Signaling Pathway and Pulmonary Fibrosis Idiopathic Signaling Pathway, are activated in the liver of animals fed a NASH diet for 20 weeks compared to a control diet (B). Orange bars denote activated pathways (positive z-score > 2), whereas blue bars denote inhibited pathways (negative z-score <- 2.0). Putative upstream regulators of the DEGs at this time point include fibrosis-related molecules such as TGFB1 and AGT (C). In the liver of mice fed a NASH diet, the gene set for collagen fibril organization is significantly enriched (D). Genes related to fibrosis, including *Aebp1*, *Col1a1*, *Col1a2*, *Mmp2*, *Mmp13*, and *Timp1*, are significantly upregulated in the liver of mice fed the NASH diet for 20 weeks, compared to the control diet (E). Statistical significance was determined by t-test, with p < 0.05 considered significant. * p < 0.05, ** p <0.01.

### Activated inflammasome response in the liver of mice fed the NASH diet for 30 weeks

At 30 weeks, there were a total of 4,132 DEGs (2,320 upregulated and 1,812 downregulated) in the liver of mice fed the NASH diet compared to mice fed a control diet. The top 40 DEGs at 30 weeks included genes related to metabolic pathways, cytochrome P450 activity, cytokine activity, and genes encoding glycoproteins **(Figure 6A)**. Using IPA, we determined that DEGs at this time point were enriched for fibrosis- and immune-related pathways, including activation of the Pulmonary Fibrosis Idiopathic Signaling Pathway, Hepatic Fibrosis Signaling Pathway, and IL-8 Signaling **(Figure 6B)**. Similar to what was observed at 10 and 20 weeks, potential upstream regulators of DEGs at 30 weeks included fibrosis- and immune-related genes, such as *TGFB-1* and *TNF* **(Figure 6C)**. Using GSEA, we determined that the gene signature identified in the liver from mice fed the NASH diet for 30 weeks was enriched for genes related to the GO term “Cytokine Secretion” **(Figure 6D)**. These cytokine secretion-related genes include genes in the inflammasome signaling pathway. Therefore, we validated a subset of inflammasome-related genes in the larger cohort of animals. We observed an increase in *Aim2*, *Casp1*, and *Nrlp3* expression in the liver of mice fed the NASH diet for 30 weeks compared to the liver of mice fed a control diet that was animal-dependent (did not reach statistical significance) **(Figure 6E)**. We also observed increased *Pycard*, *Tlr4*, and *Tlr9* expression in the liver of mice fed the NASH diet for 30 weeks **(Figure 6E)**.

**Figure 6.**
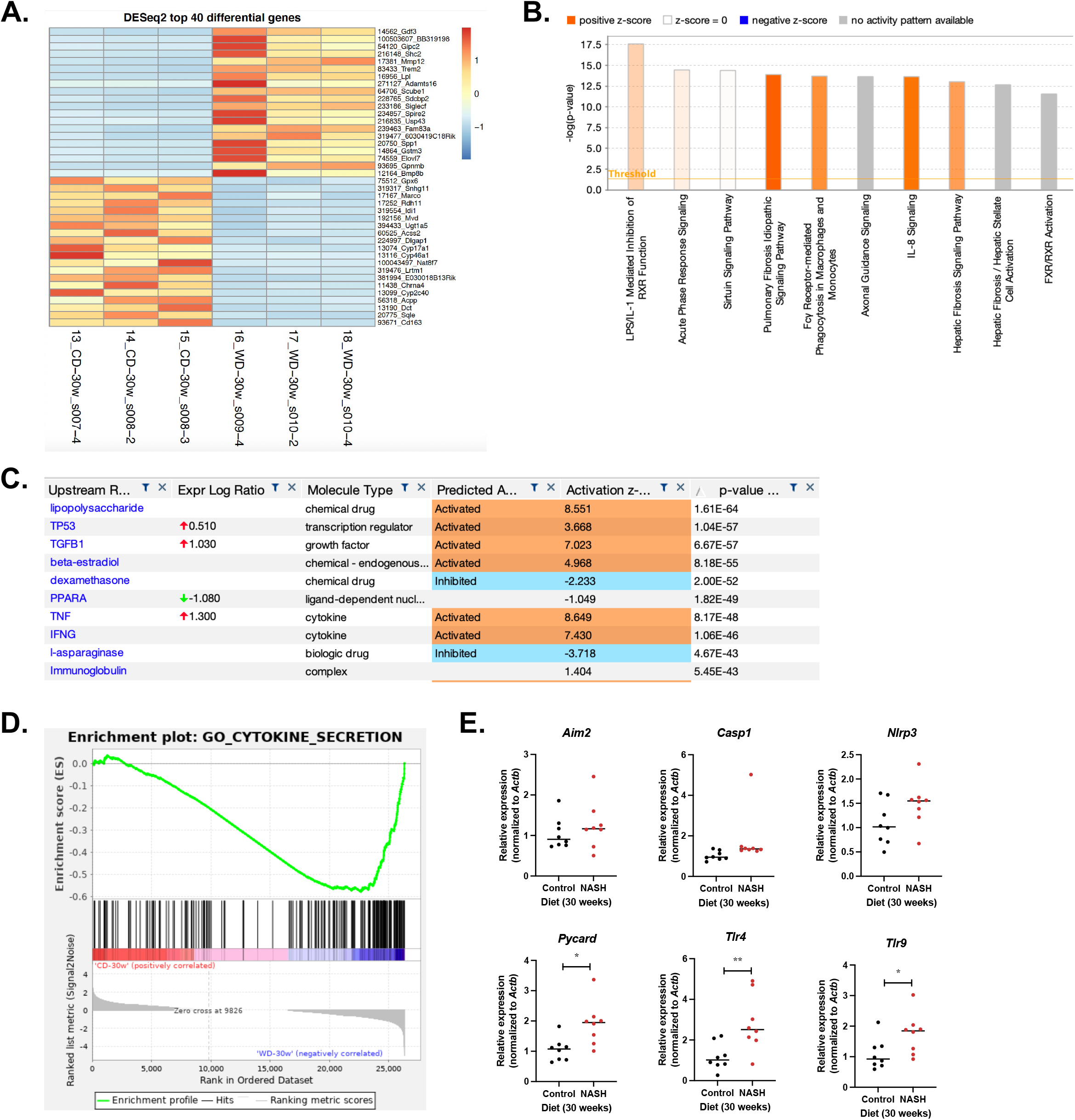
Hepatic transcriptome from mice fed a NASH diet for 30 weeks. Effect of feeding male mice either a control diet (10% kcal from fat) or NASH diet (40% kcal from fat, 20% kcal fructose, and 2% cholesterol) for 30 weeks on hepatic gene expression. Heat map of the top 40 differentially expressed genes (DEGs; A). Pathways related to fibrosis and immune response, including Hepatic Fibrosis Signaling Pathway, Pulmonary Fibrosis Idiopathic Signaling Pathway, and IL-8 signaling, are activated in the liver of animals fed a Western-style diet for 30 weeks compared to a control diet (B). Orange bars denote activated pathways (positive z-score > 2), whereas blue bars denote inhibited pathways (negative z-score <- 2.0). Putative upstream regulators of the DEGs at this time point include fibrosis- and immune-related molecules such as TGFB1 and TNF (C). In the liver of mice fed a NASH diet, the gene set for cytokine secretion is significantly enriched (D), including genes in the inflammasome pathway. Genes related to the inflammasome, including *Aim2*, *Casp1*, and *Nlrp3*, are upregulated in an animal-dependent manner. Other inflammasome-related genes, including *Pycard*, *Tlr4*, and *Tlr9*, are significantly upregulated in the liver of mice fed the NASH diet for 30 weeks compared to the control diet (E). Statistical significance was determined by t-test, with p < 0.05 considered significant. * p < 0.05, ** p <0.01.

### Longitudinal analysis of DEGs reveals early- and late-stage pathways associated with progressive liver disease

By comparing the RNA signatures at 10, 20, and 30 weeks we identified DEGs and pathways associated with early- and late-stage liver disease in the context of the NASH diet. For upregulated DEGs, 790 genes were common to all three time points (37, 36, and 34% of upregulated DEGs at 10, 20, and 30 weeks, respectively) **(Figure 7A)**. A total of 775, 604, and 775 genes were upregulated only at 10 (36%), 20 (26%), or 30 (33%), respectively **(Figure 7A)**. For downregulated DEGs, 321 genes were common to all three time points (25, 18, and 18% of downregulated DEGs at 10, 20, and 30 weeks, respectively) **(Figure 7B)**. A total of 601, 697, and 671 genes were downregulated only at 10 (47%), 20 (39%), or 30 (37%), respectively **(Figure 7B)**. Using IPA, we determined that pathways common to all three time points include pathways related to biosynthesis, disease-specific pathways, cellular immune response, neurotransmitters and other nervous system signaling, and cytokine signaling, among others **(Figure 7C)**. Specifically, we observed predicted inhibition of the Superpathway of Cholesterol Biosynthesis, and predicted activation of the Pathogen Induced Cytokine Storm Signaling, Neuroinflammation Signaling, Role of Osteoclasts in Rheumatoid Arthritis Signaling, IL-8 Signaling, TREM1 Signaling, and Hepatic Fibrosis Signaling pathways. These data suggest that immune signaling and inflammation are disrupted early in NAFLD/NASH development and continue to be drivers throughout disease progression.

**Figure 7.**
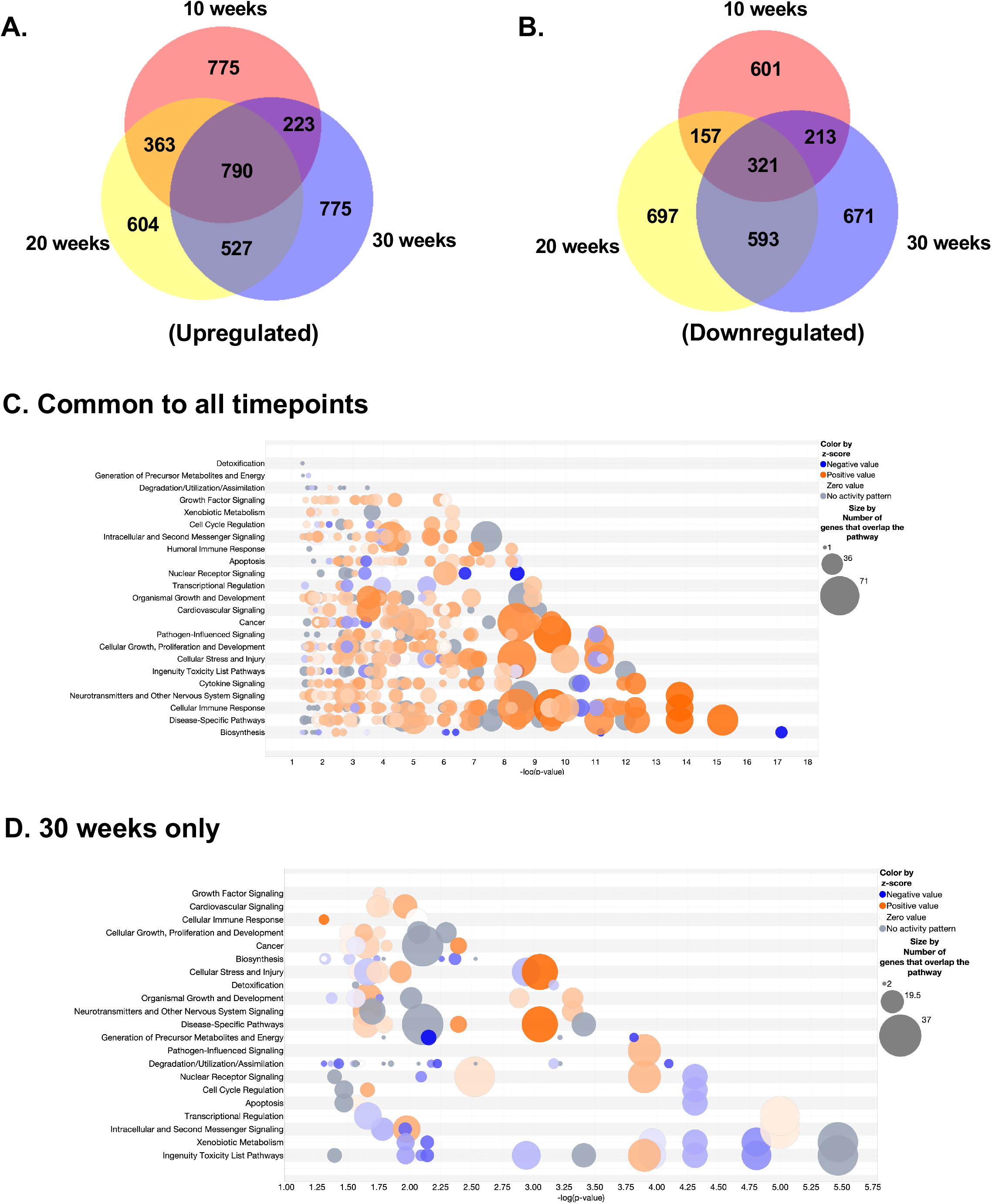
Longitudinal transcriptomic analysis from mice fed a NASH diet for up to 30 weeks. Venn diagrams report the numbers of both up- and down-regulated genes (A and B, respectively) in mice fed a NASH diet or a control diet for up to 30 weeks. Pathway analysis of differentially expressed genes that are common to all time points or specific to the 30-week time point alone are also reported (C and D, respectively).

We also identified pathways associated with the phenotype observed in the mice fed the NASH diet for 30 weeks. Using IPA, we determined that pathways altered at 30 weeks include those related to xenobiotic metabolism, intracellular and second messenger signaling, transcriptional regulation, cell cycle regulation, and nuclear receptor signaling **(Figure 7D)**. Specifically, we observed predicted inhibition of the Xenobiotic Metabolism PXR Signaling, Aryl Hydrocarbon Receptor Signaling, and Xenobiotic Metabolism CAR Signaling pathways, and predicted activation of the Sirtuin Signaling, LPS/IL-1 Mediated Inhibition of RXR Function, and Estrogen Receptor Signaling pathways. These data suggest that inhibition of xenobiotic metabolism and PXR/CAR signaling is a late-stage driver of NAFLD/NASH progression.

### Altered microbiome composition in mice fed the NASH diet

Studies have reported gut microbiota alterations in both animal models of and humans with NAFLD (12). Therefore, we performed microbiome analysis on fecal pellets collected from the mice fed the NASH diet or a control diet for 10, 20, or 30 weeks. We observed a decrease in alpha diversity (within-sample diversity) for the mice fed a NASH diet compared to those fed a control diet for 10 weeks **(Supplemental Figure 5A)**. Beta diversity (between sample diversity) was visualized using Principle Coordinates Analysis (PCoA). We identified that the bacterial composition of these samples clustered according to diet (**Supplemental Figure 5B)**. In the 20-week cohort, *Bacteroides theaiotamicron*, *Kineothrix sp000403275*, and *Lactococcus lactis* were increased, while *Kineothrix MGBC162921* was decreased after 10 weeks on the NASH diet compared to the control diet **(Supplemental Figure 5C and D)**

Decreased alpha diversity was also detected in the mice fed the NASH diet for 20 weeks compared to the control diet **(Figure 8A)**. Similar to what we observed in this cohort at 10 weeks, the bacterial composition of these samples clustered according to diet **(Figure 8B)**. The observed changes in *Bacteroides theaiotomicron* and *Lactococcus lactis* at 10 weeks persisted after 20 weeks on the NASH diet compared to the control diet **(Figure 8C and D)**. We also observed increased *Akkermansia muciniphila* and *Eubacterium MGBC101131*, as well as decreased *Bifidobacterium globosum, Faecalibacterium rodentium*, and *Muribaculum sp002358615* in these same mice **(Figure 8C and** **D**). Functional analysis of the whole genome shotgun metagenome sequencing data was performed to determine potential physiological consequences associated with the microbiome composition within each sample **(Figure 8E)**. We observed increased abundance of bacterial species associated with integral components of membrane, DNA binding, and ATP binding as well as decreased abundance of bacterial species associated with structural constituents of the ribosome and metal ion binding in mice fed the NASH diet compared to mice fed the control diet for 20 weeks **(Figure 8E)**.

**Figure 8.**
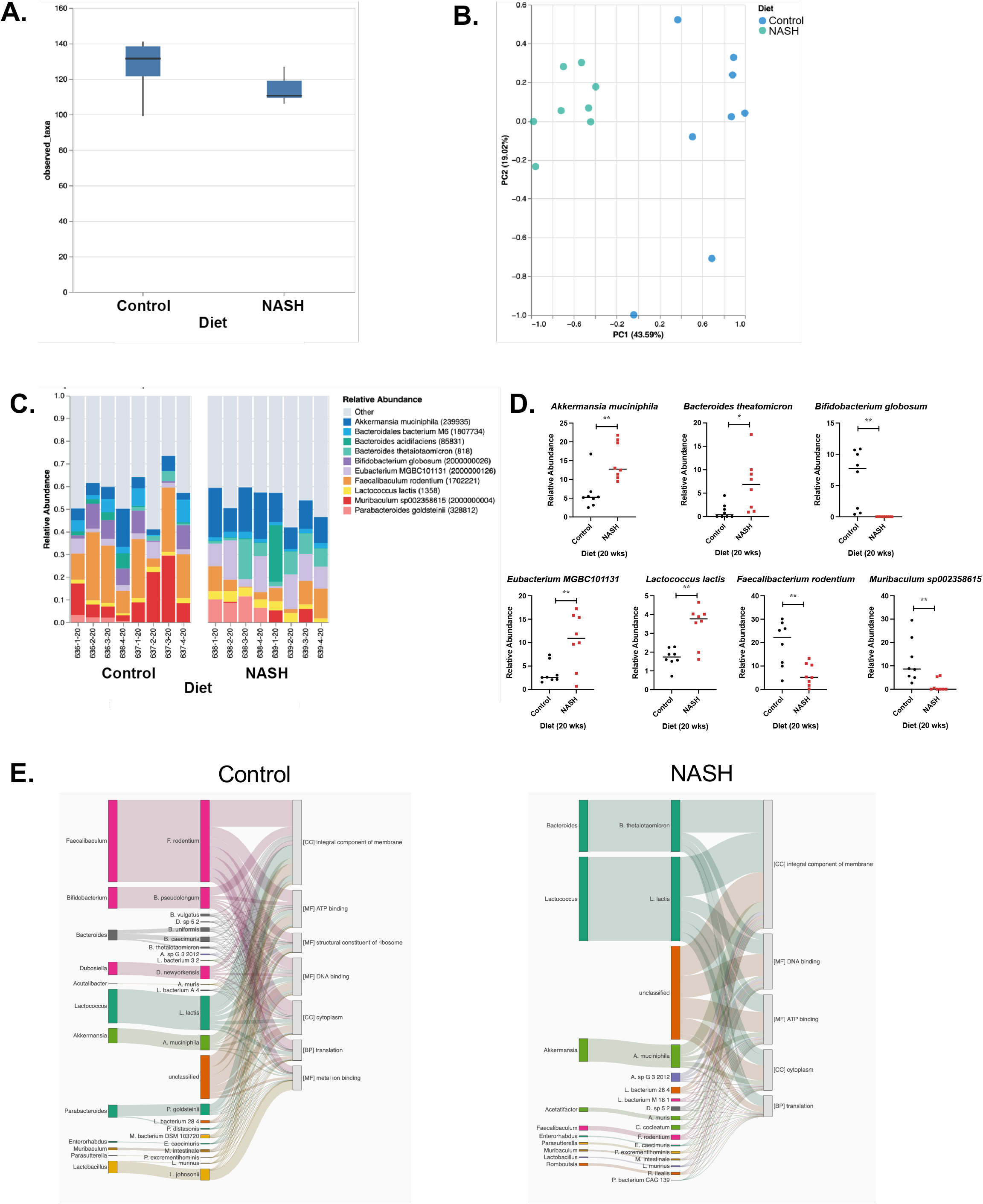
Microbiome analysis from mice fed a NASH diet for up to 20 weeks. Effect of feeding mice a NASH diet or a control diet for up to 20 weeks on beta diversity of the microbiome (A). PC1 accounts for 43.12% of the variance in beta diversity and PC2 accounts for 15.48% of the variance in beta diversity due to diet (B). Plot of relative abundance of bacterial species in the 20-week cohort after feeding a NASH or control diet for 20 weeks (C); selected species are exhibited in D. Functional impact of microbiome cellular processes (E). Statistical significance was determined by t-test, with p < 0.05 considered significant. * p < 0.05, ** p <0.01, **** p< 0.0001.

No difference in alpha diversity was observed in the 30-week cohort fed the NASH diet or control diet for 20 weeks **(Supplemental Figure 6A)**. The bacterial composition of these samples clustered according to diet **(Supplemental Figure 6B)**. In the 30-week cohort, we observed increased *Akkermansia muciniphila, Alistipes MGBC111863*, *Bacteroides acidifaciens*, and a trend toward increased *Paramuribaculum intestinale*, as well as decreased *Alistipes sp002428825, Dubosiella newyorkensis*, and *Porphyromonodaceae bacterium UBA7207* after 20 weeks on the NASH diet compared to the control diet **(Supplemental Figure 6C and D)**. The observed change in *Akkermansia muciniphila* is consistent with alterations observed in the 20-week cohort at the same time point **(Figure 8C and D)**.

Decreased alpha diversity was observed in the mice fed the NASH diet for 30 weeks compared to the control diet **(Figure 9A)**. As expected, the bacterial composition of these samples clustered according to diet **(Figure 9B)**. The changes in *Akkermansia muciniphila, Alistipes sp002428825*, and *Bacteroides acidifaciens* observed in this cohort at the 20-week timepoint persisted after 30 weeks on the NASH diet compared to the control diet **(Figure 9C and D)**. We also observed increased *Bacteroides sp002491635* and *Kineothrix sp000403275*, as well as decreased *Faecalibacterium rodentium* in these same mice **(Figure 9C and D)**. Functional analysis revealed increased abundance of bacterial species associated with integral components of membrane and metal ion binding, as well as a decreased abundance of bacterial species associated with structural constituents of the ribosome, ribosome, and translation in mice fed the NASH diet compared to mice fed the control diet for 30 weeks **(Figure 9E)**.

**Figure 9.**
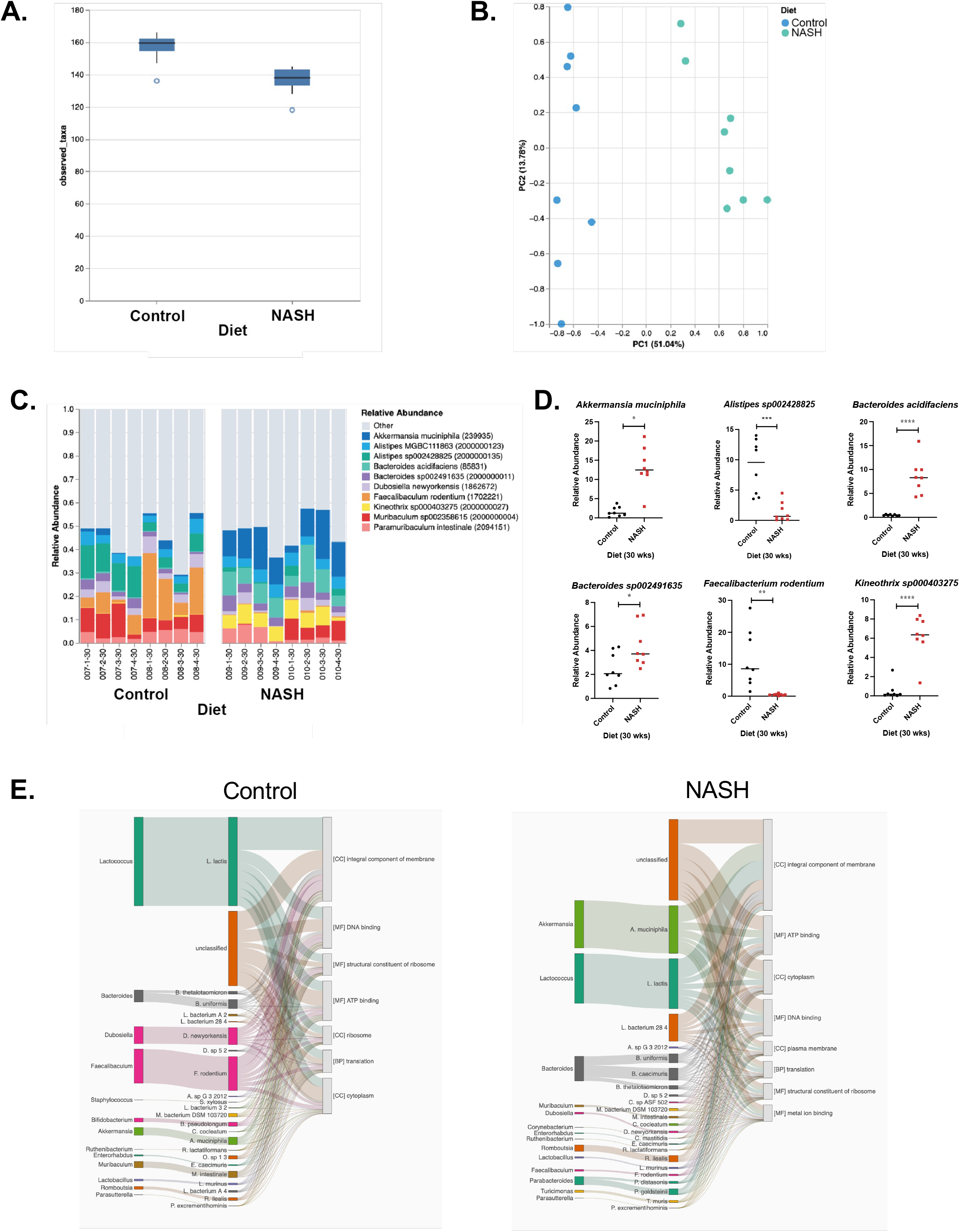
Microbiome analysis from mice fed a NASH diet for up to 30 weeks. Effect of feeding mice a NASH diet or a control diet for up to 30 weeks on beta diversity of the microbiome (A). PC1 accounts for 31.68% of the variance in beta diversity, and PC2 accounts for 20.87% of the variance in beta diversity due to diet (B). Plot of relative abundance of bacterial species in the 20-week cohort after feeding a NASH or control diet for 20 weeks (C); selected species are exhibited in D. Functional impact of microbiome cellular processes (E). Statistical significance was determined by t-test, with p < 0.05 considered significant. * p < 0.05, ** p <0.01, *** p < 0.001, **** p< 0.0001.

## DISCUSSION

Although the global prevalence of NAFLD/NASH is increasing at an alarming rate, there are no approved medications to treat this disease. Therefore, there is urgency to develop new prevention and therapeutic paradigms. Preclinical models of NAFLD/NASH/HCC for the development of such paradigms have historically focused on genetic manipulation or the use of carcinogens; the use of solely nutritionally based models, which more closely reflects the causative agents for humans, has been less common. Studies have shown that a diet (GAN-DIO-NASH) high in fat (palm oil), fructose, and cholesterol can promote the development of NAFLD/NASH/HCC in mice (13-15). Whereas these studies have supported the use of nutritionally based models of NAFLD/NASH/HCC for preclinical studies, they were conducted in *ob/ob* mice and did not comprehensively profile phenotypic and transcriptomic changes over time. Therefore, we set out to perform a longitudinal analysis of the effects of the NASH diet, in the absence of gross hyperphagia, to further support the use of this exclusively dietary model for preclinical studies of prevention and therapeutic paradigms.

We fed C57BL/6J mice the NASH diet for 10, 20, and 30 weeks and performed biochemical, histopathological, transcriptomic, and microbiome analyses at each time point. We chose C57BL/6J mice over *ob/ob* mice due to limitations in the translational utility of *ob/ob* mice as they are resistant to fibrosis, do not develop NASH, and do not develop HCC (16-18). Like previous reports utilizing this diet, we observed progressive worsening of NAFLD/NASH over time based on liver histopathological and biochemical analysis. In addition, mice fed the NASH diet exhibited increased body weight and fat mass, most likely due to modest increases in food intake during the first few months on the diet, as well as the addition of fat, cholesterol, and fructose to the diet. Activity levels were not influenced by the diet. Interestingly, caloric intake was not different between the two groups at the 20- and 30-week time points, suggesting that the content of the diet itself and/or early excess calories triggered the subsequent changes in the microbiome as well as hepatic transcriptome and metabolism. In support of the idea that the early exposure to the diet profoundly alters liver physiology, the livers of the NASH diet-fed animals markedly switch their preference for storing carbons (i.e., decreased hepatic glycogen and increased hepatic triglycerides in the NASH compared to control mice), and have early, but subclinical (i.e., absence of elevated serum ALT or hyperlipidemia), signs of hepatic steatosis, and fibrosis.

Longitudinal analysis of liver DEGs identified early- and late-stage drivers of NAFLD/NASH development and progression. Immune-related genes are early-stage drivers of NAFLD/NASH, as evidenced by the enrichment of adaptive immune response genes in the gene signature after 10 weeks on the NASH diet. Whereas it has been recognized that innate immune mechanisms play a key role in promoting inflammation in NASH, increasing evidence supports a role for adaptive immunity in promoting liver inflammation as well (19). We identified that dysregulation of immune signaling persists at 20 and 30 weeks on the NASH diet. Activated (predicted) pathways that are common to all three time points include Pathogen Induced Cytokine Storm Signaling, Neuroinflammation Signaling, IL-8 Signaling, and TREM1 Signaling, among others. These results highlight the importance of systemic inflammation due to disrupted immune signaling in the initial development, and worsening, of NAFLD. Consistent with our results, another study reported altered innate and adaptive immune signaling in the liver of C57BL/6J mice fed the NASH diet for 38-44 weeks (14),.

We utilized IPA to identify late-stage drivers of NAFLD/NASH development/progression defined as the liver DEGs that were only present in the gene signature after 30 weeks on the NASH diet. Interestingly, these DEGs were enriched for genes related to xenobiotic metabolism signaling. Specifically, Xenobiotic Metabolism PXR Signaling, Xenobiotic Metabolism Constitutive Androstane Receptor (CAR) Signaling, and Aryl Hydrocarbon Signaling pathways are predicted to be inhibited. Furthermore, the LPS/IL-Mediated Inhibition of RXR Function pathway is expected to be activated. Collectively, these results suggest that inhibition of nuclear receptor signaling pathways related to xenobiotic metabolism play an important role in the later stages of NAFLD/NASH development. Activation of the CAR appears to ameliorate diet-induced hepatic steatosis (20) and inflammation (21) in mice. The effects of CAR activation on the development/progression of hepatic fibrosis and hepatocarcinogenesis are still not entirely clear (22). The role of activation of the pregnane X receptor (PXR) in the development/progression of NAFLD/NASH is complex (22). In a mouse model of diet-induced obesity, PXR knockout reduced hepatic steatosis and insulin resistance (23). The same PXR deletion strategy was introduced into *ob/ob* mice, increasing metabolic expenditure and decreasing gluconeogenesis (23). In a different PXR knockout model, PXR-KO mice gained less weight when fed a high fat diet. However, these mice exhibited hepatomegaly, hyperinsulinemia, and hyperleptinemia (24). PXR activation is generally anti-inflammatory and has also been shown to attenuate chemically induced liver fibrosis in mice (25). The role of aryl hydrocarbon receptor (AhR) activation in diet induced NAFLD development is also not clear. In one study, liver specific knockout of AhR accelerated the development of hepatic steatosis and promoted hepatic inflammation in mice fed a high fat diet (26). However, in another study, overexpression of AhR induced hepatic steatosis in mice fed a high fat diet (27). Interestingly, despite their exacerbated hepatosteatosis, these mice were protected from diet-induced obesity and insulin resistance. Follow-up studies to elucidate the impact of inhibition of CAR/PXR/AhR signaling in our model of NAFLD/NASH are beyond the scope of this paper; more studies examining the roles of CAR/PXR/AhR in diet induced NAFLD/NASH/HCC progression are certainly warranted.

The gut microbiome is thought to play a role in the development of NAFLD/NASH. Therefore, we also performed a longitudinal analysis of the gut microbiome composition in mice fed the NASH diet for up to 20 (10-and 20-week timepoints) or 30 (20- and 30-week timepoints) weeks. An increased abundance of *Bacteroides* was observed in mice fed the NASH diet, starting at 10 weeks and remaining increased at 20 and 30 weeks. Importantly, elevated levels of *Bacteroides* have been reported in patients with NAFLD/NASH (28-32), and has also been associated with HCC in patients with cirrhosis and NAFLD (33). In our study, mice fed the NASH diet for 20 and 30 weeks had an increased abundance of *Akkermansia muciniphila*. However, this result is inconsistent with what has been reported in patients with NAFLD. For example, decreased *Akkermansia muciniphila* has been reported in obese patients with NAFLD (34), in patients with type 2 diabetes and NAFLD (34), in patients with NAFLD with increased liver enzymes (35), and in patients with biopsy-confirmed NASH (36). Children with NAFLD/NASH exhibit similar changes (37-40). Studies have suggested a protective role for *Akkermansia muciniphila* in the development of NAFLD/NASH, with probiotic administration preventing the development of and ameliorating complications related to the disease (41). Thus, the observed increase in our study may represent a compensatory mechanism being activated in the mouse. However, the increased abundance in *Akkermansia muciniphila* was not sufficient to prevent the progression of NAFLD/NASH in the mice fed the NASH diet for 20 weeks and beyond. We also observed a decreased abundance of *Porphyromonadaceae* after feeding the 30-week cohort mice the NASH diet for 20 weeks. A previous report also observed decreased abundance of *Porphyromonadaceae* using this same diet in *ob/ob* mice (13) and in patients with NAFLD compared to healthy controls (42). Given that gut dysbiosis occurs during NAFLD development and progression in preclinical models and in patients, the administration of probiotics, parabiotics, or postbiotics is one avenue of further exploration for the management of NAFLD (43).

The current mouse model phenocopies many of the features of NAFLD and NASH in humans, including but not limited to elevated serum ALT, hypercholesterolemia, the profound steatosis, fibrosis, and selected changes in the microbiome. Nevertheless, there are some notable differences, particularly in regards to the metabolic aspects of MAFLD and NAFLD in relation to the common comorbidity of metabolic syndrome. For example, we were unable to detect profound signatures of insulin resistance, as the NASH diet did not promote hyperinsulinemia or provoke changes in insulin tolerance. However, insulin tolerance testing is less sensitive than the gold-standard hyperinsulinemic-euglycemic clamp (44-46), and it should be noted that changes in hepatic insulin sensitivity could be revealed by the tissue-specific nature of a clamp study. Similarly, changes in oral glucose tolerance, which reflects a composite of beta cell function and glucose disposal along with the incretin response, were not detected, which is surprising given that humans with NAFLD have a reduction in secretion of the incretin glucagon-like peptide 1 (GLP-1) (47). Because of the altered incretin response in human NAFLD and the availability of a variety of incretin mimetics for treating type 2 diabetes, more recent research has focused on targeting the incretin system in NASH and potentially repurposing these type 2 diabetes therapeutics for NAFLD and NASH. It is possible that altering the fasting duration or glucose load would reveal changes in glucose tolerance not observed in the present study. In addition to the technical limitations of the tolerance testing, the current dietary regimen provoked only a mild, albeit significant, increase in body weight and adiposity; perhaps more severe increases in those parameters or even extending the duration on the diet would recapitulate more of the metabolic derangements observed in human NAFLD. Finally, we did not evaluate changes in blood pressure or other cardiovascular parameters reflected by the metabolic syndrome.

NAFLD prevalence in humans is higher in males than in females during the reproductive years (48). However, postmenopausal women exhibit high rates of NAFLD, which suggest the protective effects of estrogen and other sex hormones. The current study focused exclusively on male mice given that female mice are also largely protected against metabolic dysfunction and profound obesity when exposed to high caloric diets. Nevertheless, future work should consider the impact of sex and sex hormones in animal models of NAFLD and NASH.

In summary, we have provided a comprehensive analysis of changes in the transcriptome, microbiome, and physiological, pathological, and biochemical parameters throughout a 30-week exposure to a NASH-promoting diet in male mice. This exclusively dietary NASH model recapitulates many, but not all, the features, of NAFLD, MAFLD, and NASH in humans. Specifically, we identified transcriptomic changes in the livers of NASH-diet fed animals that reflect immune response activation. We demonstrated significant overlap to changes in the microbiome in humans with NAFLD and NASH. Finally, the biochemical and pathological parameters closely mirror those of the human disease. This longitudinal analysis highlights the dynamic changes throughout the course of NAFLD and NASH development and will be useful for future intervention time points.

## ACKNOWLEDGEMENTS

The authors thank the Animal Resources Core at the City of Hope, supported by the National Cancer Institute of the NIH (P30CA033572). M.S. was supported by a National Cancer Institute Cancer Metabolism Training Program Postdoctoral Fellowship (T32CA221709). This work was supported, in part, by pilot funding from the Beckman Research Institute of City of Hope including a Junior Investigator Research Development Award (awarded to L.S.T.), City of Hope-Ludwig Cancer Research Hilton Foundation Pilot Award (awarded to L.S.T.), and Innovative Inter-programmatic Collaborations in Cancer Control and Population Sciences Award (awarded to P.T.F).

## AUTHOR STATEMENT

**Marissa Saenz:** Data curation, Investigation, Validation, Visualization, and Editing; **Jillian C. McDonough:** Data curation, Investigation, and Editing; **Elizabeth Bloom-Saldana:** Data curation, Investigation, and Editing; **Jose Irimia:** Data curation, Investigation, Validation, Visualization, and Editing; **Emily L. Cauble:** Data curation, Investigation, and Editing; **Ashly Castillo:** Data Curation, Investigation, and Editing; **Patrick T. Fueger:** Conceptualization, Methodology, Project administration, Resources, Supervision, Visualization, Writing original draft, and Editing; **Lindsey Treviño:** Conceptualization, Data Curation, Formal analysis, Methodology, Project administration, Resources, Supervision, Visualization, Writing original draft and Editing.

## LEGENDS FOR SUPPLEMENTAL DATA

**Supplemental Table 1. Metabolic testing in mice fed a NASH diet for up to 30 weeks.** Glucose and insulin tolerance testing was performed on the three cohorts at selected time points. Reported are the areas-under-the-curves (AUC) for glucose tolerance testing and areas-of-the-curves (AOC) for insulin tolerance testing. Statistical significance was tested using Student’s t-test at individual time points. No statistically significant changes were observed for these parameters based on diet at any time point.

**Supplemental Figure 1. Serum hormone and metabolic substrates from mice fed a NASH diet for up to 30 weeks.** Effect of feeding male mice with either a control diet (10% kcal from fat) or a NASH diet (40% kcal from fat, 20% kcal fructose, and 2% cholesterol) on serum levels of albumin (A), insulin (B), free glycerol (C), triglycerides (D), and non-esterified fatty acids (NEFAs; E). Statistical significance was tested using Student’s t-test at individual time points. No statistically significant changes were observed for these parameters based on diet at any time point.

**Supplemental Figure 2. Metabolic caging in mice fed a NASH diet for 20 weeks.** Instrumented metabolic caging was used to quantify food consumption (A-C), water consumption (D-F), energy expenditure (G-I), net energy balance (J-L), locomotor activity (M-O), and respiratory exchange ratio (P-Q) over a three-day period in mice fed a control or NASH diet for 10 weeks. n = 8; * p < 0.05.

**Supplemental Figure 3. Metabolic caging in mice fed a NASH diet for 30 weeks.** Instrumented metabolic caging was used to quantify food consumption (A-C), water consumption (D-F), energy expenditure (G-I), net energy balance (J-L), locomotor activity (M-O), and respiratory exchange ratio (P-Q) over a three-day period in mice fed a control or NASH diet for 10 weeks. n = 8; * p < 0.05.

**Supplemental Figure 4. PCA analysis of microbiome from mice fed a NASH diet.** Principal Component Analysis (PCA) summarizing the expression values for each sample in the 2D plane of PC1 and PC2 for diet (A) and both diet and timepoint (B). PC1 accounts for the most amount of variation across the samples, 61.86% for diet and 52.0% for diet and timepoint. PC2 accounts for the second most variation across samples, 13.1% for diet and 19.11% for diet and time point.

**Supplemental Figure 5. Microbiome analysis from mice fed a NASH diet for 10 weeks.** Alpha diversity analysis of the microbiome in mice fed a Western-style diet or a control diet for up to 20 weeks at the 10-week time points using the Shannon, Simpson, and Observed taxa methods (A). PC1 accounts for 51.04% of the variance in beta diversity and PC2 accounts for 13.78% of the variance in beta diversity due to diet (B). Plot of relative abundance of bacterial species in the 20-week cohort after feeding a NASH or control diet for 10 weeks (C); selected species are exhibited in D. Statistical significance was determined by t-test, with p < 0.05 considered significant. * p < 0.05, ** p <0.01.

**Supplemental Figure 6. Microbiome analysis from mice fed a NASH diet for 20 weeks.** Alpha diversity analysis of the microbiome in mice fed a Western-style diet or a control diet for up to 30 weeks at the 20-week time points using the Shannon, Simpson, and Observed taxa methods (A). PC1 accounts for 46.09% of the variance in beta diversity and PC2 accounts for 18.29% of the variance in beta diversity due to diet (B). Plot of relative abundance of bacterial species in the 30-week cohort after feeding a NASH or control diet for 20 weeks (C); selected species are exhibited in D. Statistical significance was determined by t-test, with p < 0.05 considered significant. * p < 0.05, ** p <0.01, **** p< 0.0001.

