## Supplementary material for "Longitudinal analysis of a dietary mouse model of non-alcoholic fatty liver disease (NAFLD) and non-alcoholic steatohepatitis (NASH)": Supplmental Data

**Supplemental Figure 6. Microbiome analysis from mice fed a NASH diet for 20 weeks.** Alpha diversity analysis of the microbiome in mice fed a Western-style diet or a control diet for up to 30 weeks at the 20-week time points using the Shannon, Simpson, and Observed taxa methods (A). PC1 accounts for 46.09% of the variance in beta diversity and PC2 accounts for 18.29% of the variance in beta diversity due to diet (B). Plot of relative abundance of bacterial species in the 30-week cohort after feeding a NASH or control diet for 20 weeks (C); selected species are exhibited in D. Statistical significance was determined by t-test, with $p < 0.05$ considered significant. * $p < 0.05$, ** $p < 0.01$, **** $p < 0.0001$.

### Supplemental Table 1.

| | | 10 weeks | | | | 20 weeks | | | | 30 weeks | | | |
|---|---|---|---|---|---|---|---|---|---|---|---|---|---|
| | | Control | | NASH | | Control | | NASH | | Control | | NASH | |
| | Timepoint (weeks) | Mean | S.D. | Mean | S.D. | Mean | S.D. | Mean | S.D. | Mean | S.D. | Mean | S.D. |
| GTT (AUC) | 10 | 4767 | 3241 | 8587 | 3947 | 2761 | 6699 | 12256 | 6658 | 7143 | 2175 | 10379 | 1789 |
| | 20 | n.d. | n.d. | n.d. | n.d. | 7932 | 3310 | 9759 | 5107 | 6347 | 2072 | 1180 | 5018 |
| | 30 | n.d. | n.d. | n.d. | n.d. | n.d. | n.d. | n.d. | n.d. | 3803 | 3520 | 1299 | 1628 |
| ITT (AUC) | 10 | 10547 | 4439 | 11945** | 3888** | 15807 | 7011 | 27053 | 5260 | 10001** | 3351** | 11132** | 2505** |
| | 20 | n.d. | n.d. | n.d. | n.d. | 13692 | 6653 | 16238* | 3695* | 14313** | 6043** | 13144 | 2107 |
| | 30 | n.d. | n.d. | n.d. | n.d. | n.d. | n.d. | n.d. | n.d. | 8472** | 7537** | 5176 | 4893 |

* n = 6
** n = 7
n.d. = not determined

### Supplemental Figure 1.

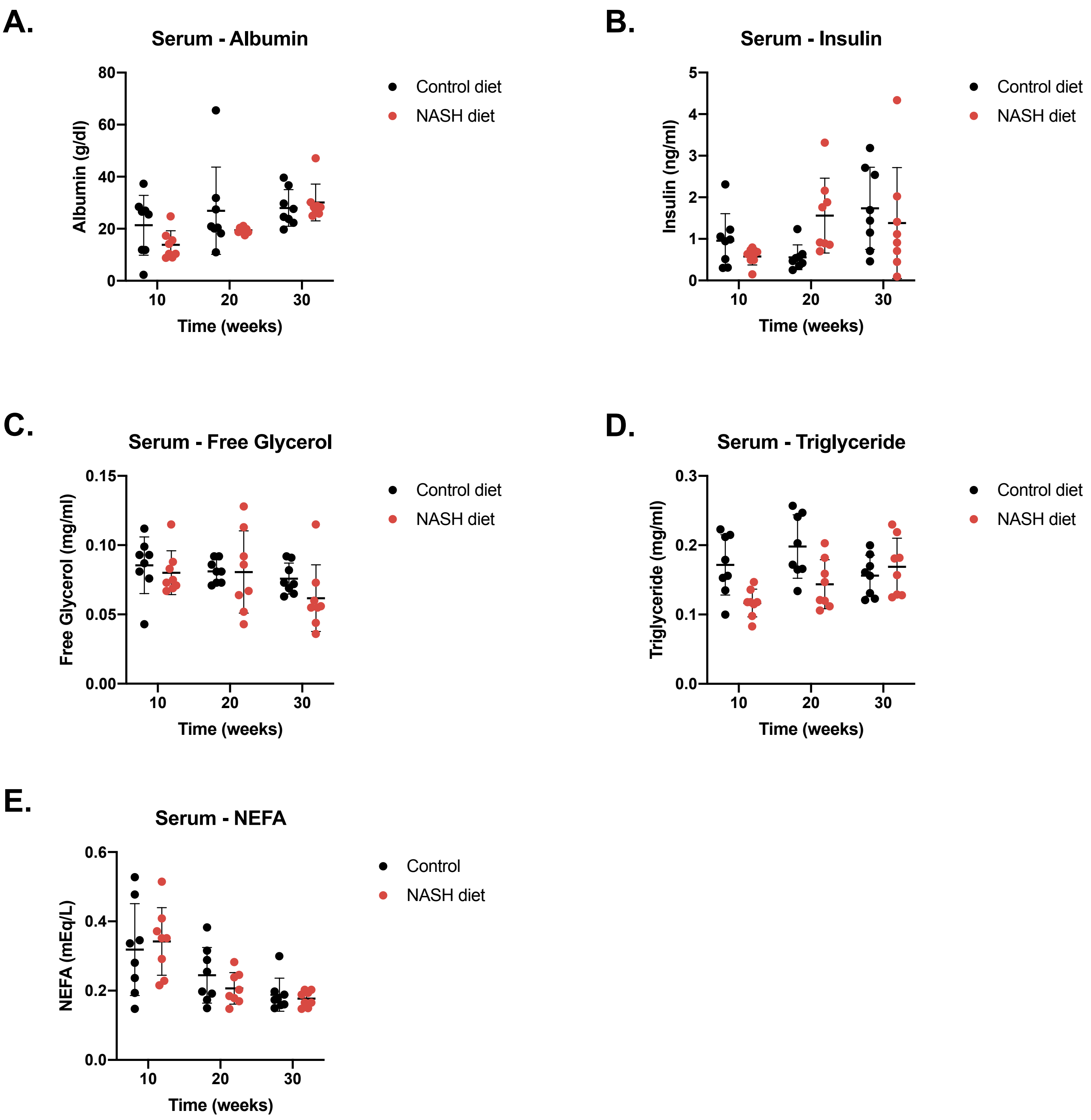

#### Supplemental Figure 2.

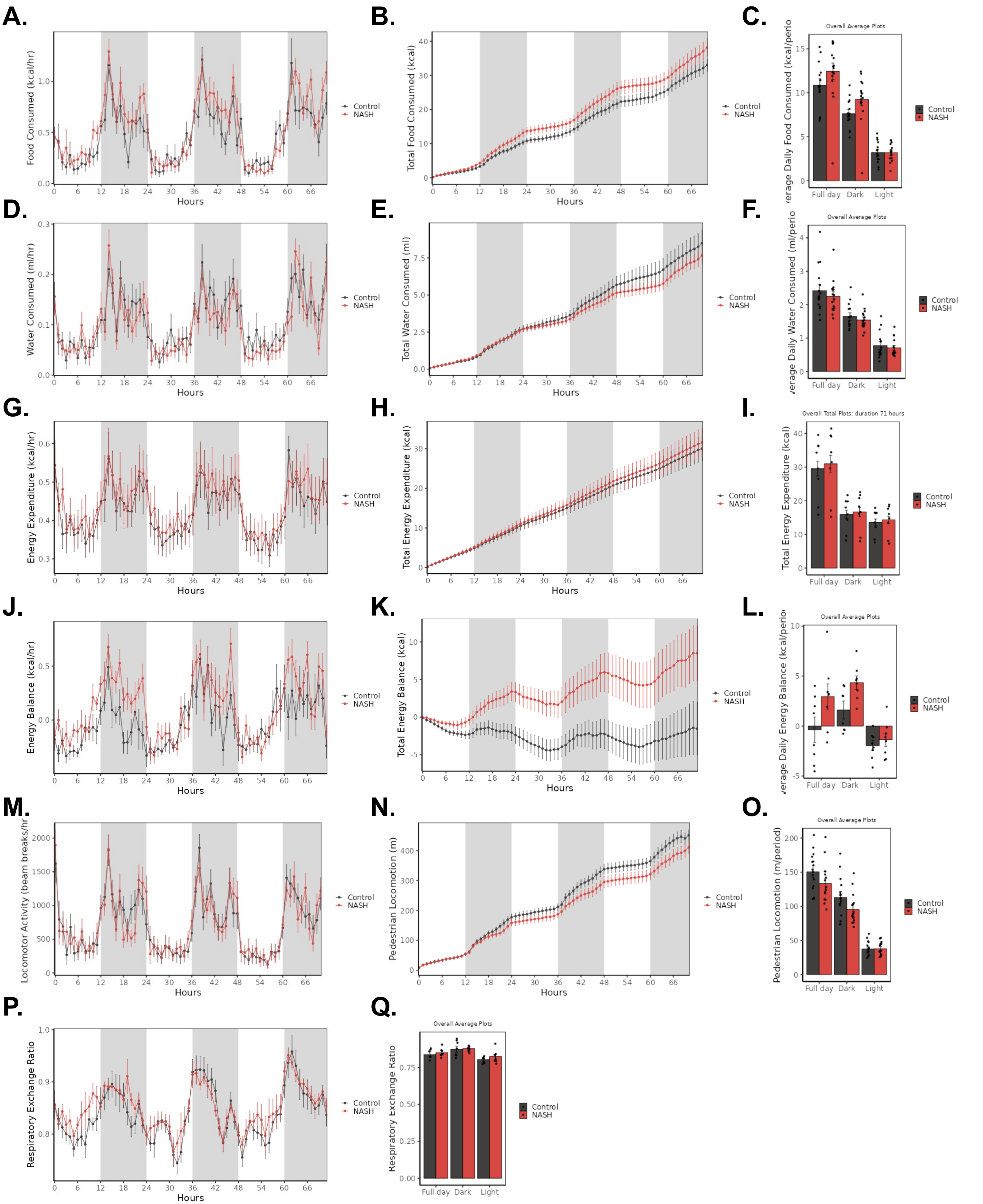

### Supplemental Figure 3.

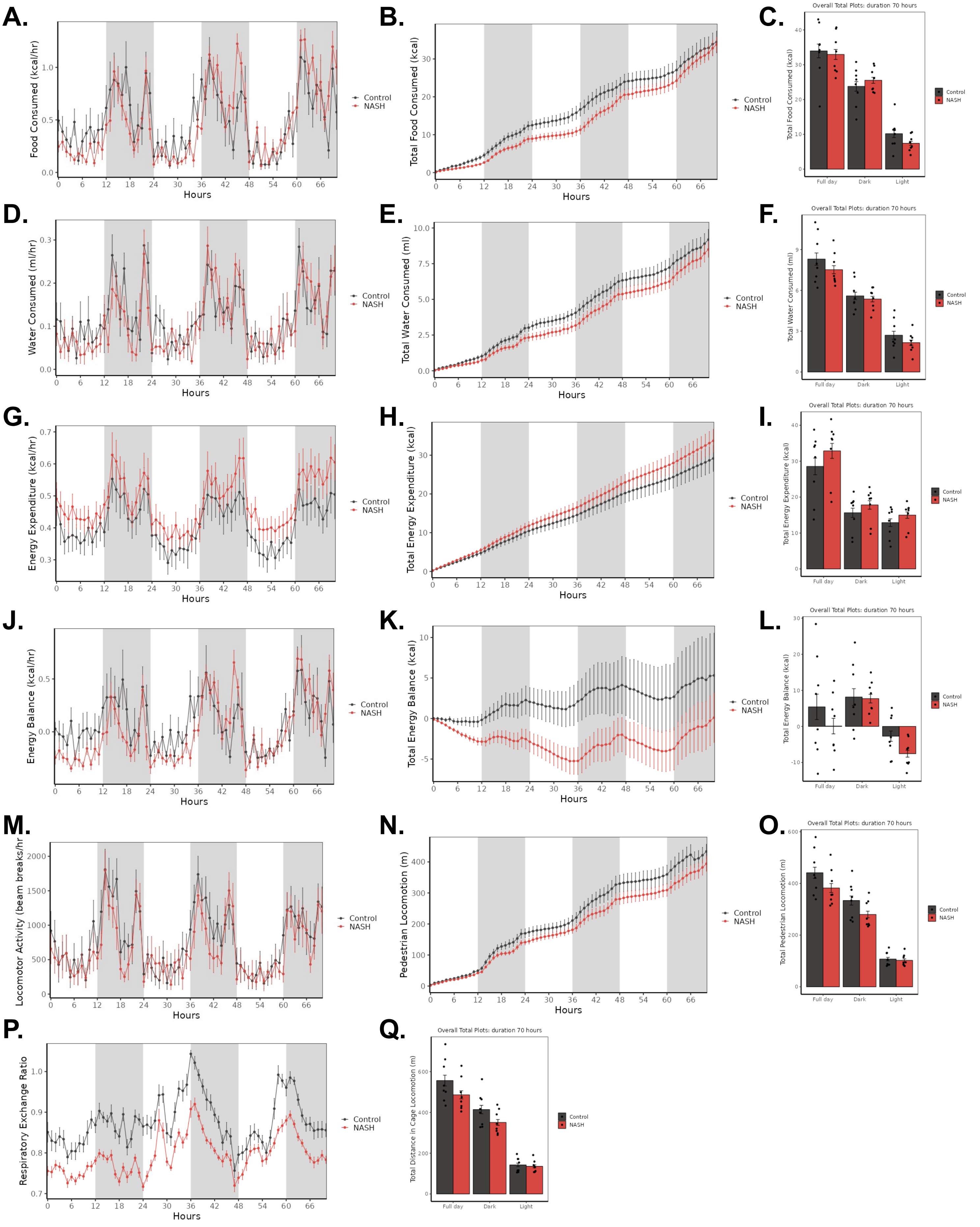

**Supplemental Figure 4.**

**A.**

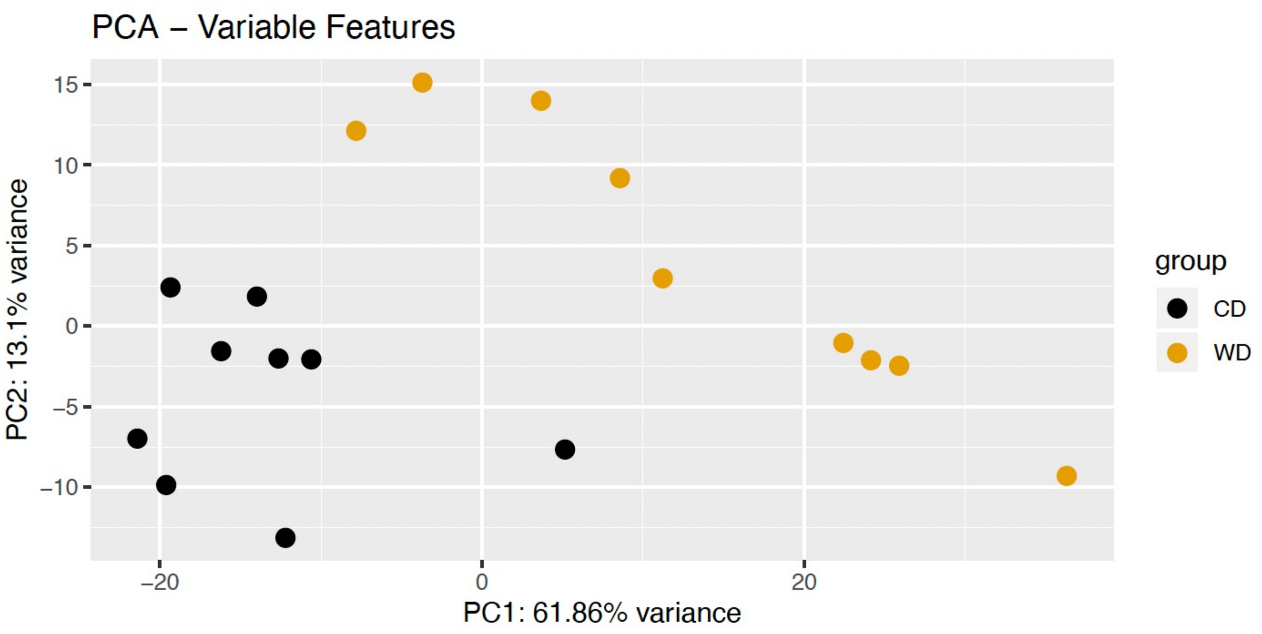


**B.**

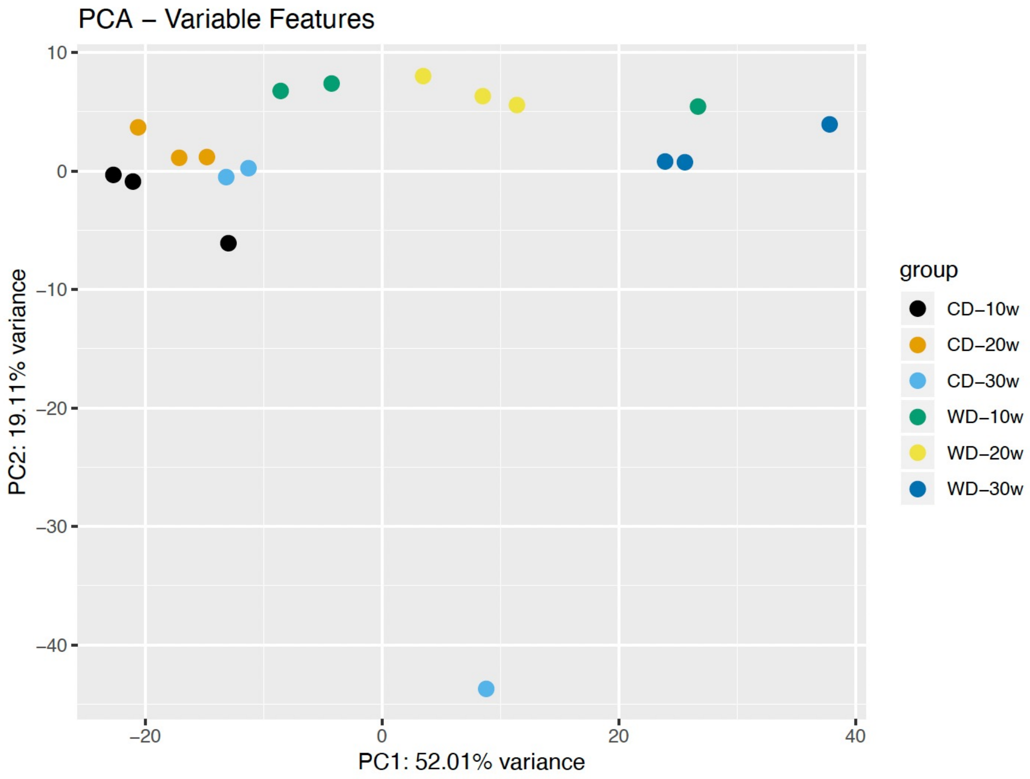

### Supplemental Figure 5.

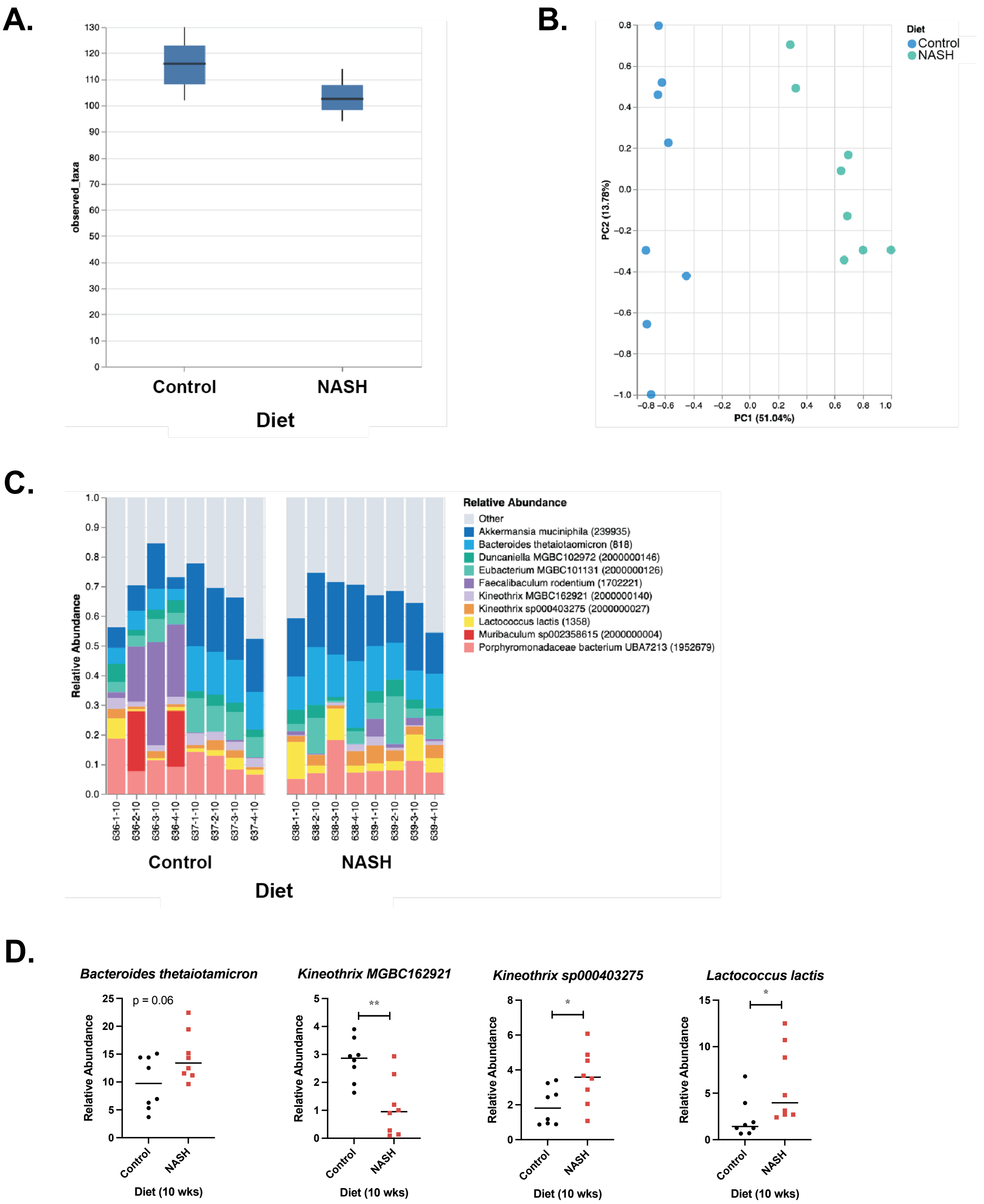

### Supplemental Figure 6.

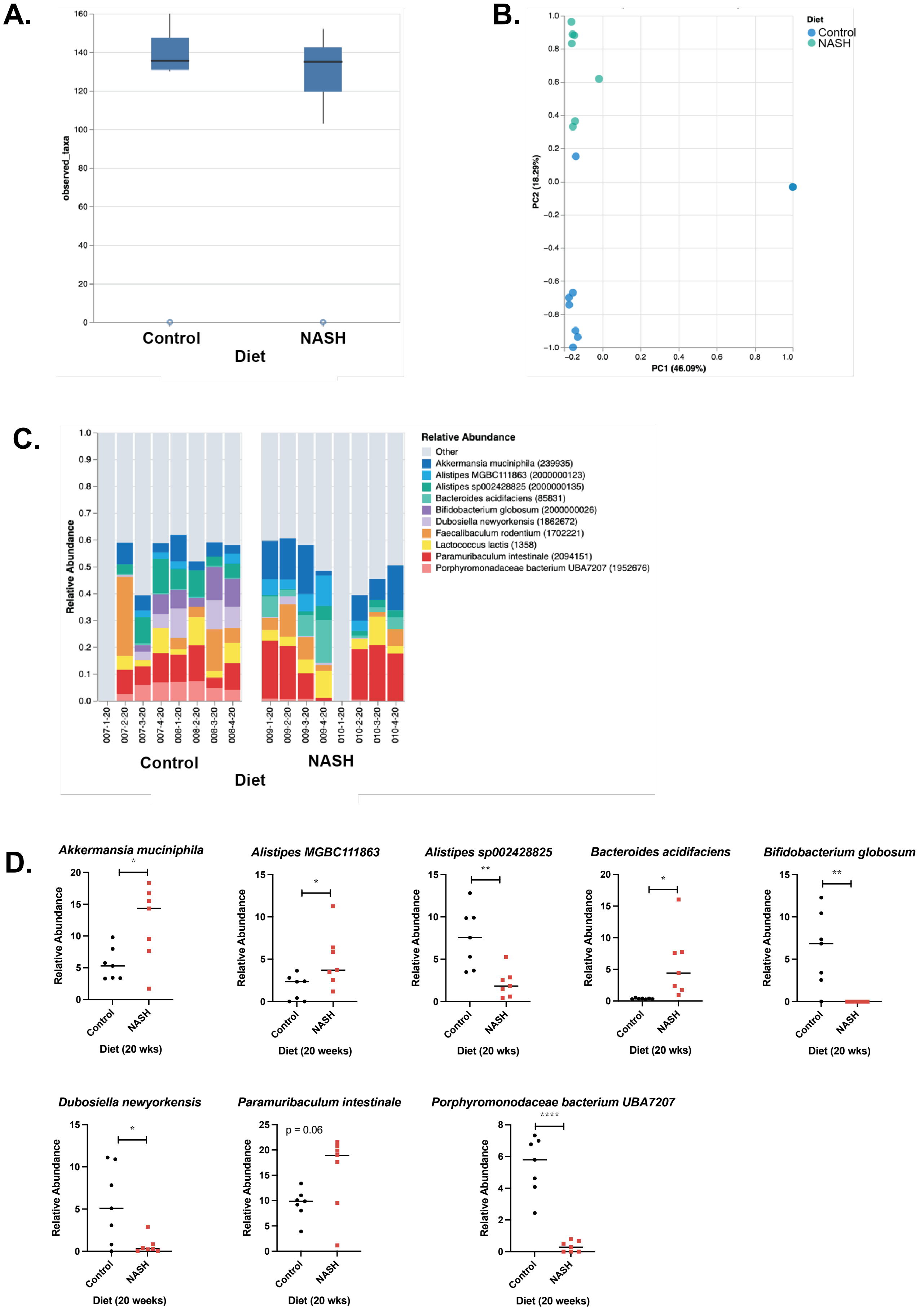
